# Brain-Specific Deletion of GIT1 Impairs Cognition and Alters Phosphorylation of Synaptic Protein Networks Implicated in Schizophrenia Susceptibility

**DOI:** 10.1101/290312

**Authors:** Daniel M. Fass, Michael C. Lewis, Rushdy Ahmad, Matthew J. Szucs, Qiangge Zhang, Morgan Fleishman, Dongqing Wang, Myung Jong Kim, Jonathan Biag, Steven A. Carr, Edward M. Scolnick, Richard T. Premont, Stephen J. Haggarty

## Abstract

Despite tremendous effort, the molecular and cellular basis of cognitive deficits in schizophrenia remain poorly understood. Recent progress in elucidating the genetic architecture of schizophrenia has highlighted the association of multiple loci and rare variants that may impact susceptibility. One key example, given their potential etiopathogenic and therapeutic relevance, is a set of genes that encode proteins that regulate excitatory glutamatergic synapses in brain. A critical next step is to delineate specifically how such genetic variation impacts synaptic plasticity and to determine if and how the encoded proteins interact biochemically with one another to control cognitive function in a convergent manner. Towards this goal, here we study the roles of GPCR-kinase interacting protein 1 (GIT1), a synaptic scaffolding and signaling protein with damaging coding variants found in schizophrenia patients, as well as copy number variants found in patients with neurodevelopmental disorders. We generated conditional neural-selective *GIT1* knockout mice and find that these mice have deficits in fear conditioning learning and spatial memory. Using global quantitative phospho-proteomics, we revealed that GIT1 deletion in brain perturbs specific networks of GIT1-interacting synaptic proteins. Importantly, several schizophrenia and neurodevelopmental disorder risk genes are present within these networks. We propose that GIT1 regulates the phosphorylation of a network of synaptic proteins and other critical regulators of neuroplasticity, and that perturbation of these networks may contribute to cognitive deficits observed in schizophrenia and neurodevelopmental disorders.

## Introduction

Neuropsychiatric disorders are devastating illnesses affecting brain function that cause tremendous suffering in patients and families worldwide. Intellectual disability neurodevelopmental (ID/NDD) disorders, with an overall prevalence of ∼1%, comprise a heterogeneous group of largely monogenic disorders that feature severe cognitive deficits either alone or in combination with syndromes of effects across other body systems (Vissers et al., 2016). Schizophrenia (SCZ), with a lifetime prevalence of ∼0.7% (McGrath et al., 2008), features positive symptoms (hallucinations, delusions), negative symptoms (flat affect, avolition), and cognitive deficits (Kahn & Keefe, 2013; Tandon et al., 2013). While ID/NDD and SCZ are clearly distinct disorders (e.g. ID/NDD is typically found in young children, while SCZ typically first presents in late adolescence/early adulthood), the shared domain of cognitive deficits suggests some convergent dysfunction.

Recent large-scale exome sequencing and genome-wide association efforts have yielded tremendous progress in identifying the genetic risk factors for neuropsychiatric diseases. For ID/NDD, over 700 risk genes have been identified (Vissers et al., 2016). Common and rare variants contributing to schizophrenia (SCZ) risk have been identified at over 100 genomic loci (Ripke et al., 2014; Fromer et al., 2014; Purcell et al., 2014; Genovese et al., 2016; Pardiñas et al., 2018). In comparisons of the cell biological functions of ID/NDD and SCZ risk genes, a number of common themes emerge (Cristino et al., 2014; Hormozdiari et al., 2015), which may reflect the commonality of cognitive deficits across these disorders. In particular, numerous ID/NDD and SCZ risk genes play key roles within pre-synaptic and post-synaptic specializations in neurons in the central nervous system. On the pre-synaptic side, studies suggest dysregulation of presynaptic vesicle recruitment, docking, release, and recycling in both ID/NDD (e.g. Broek et al., 2016; Pechstein et al., 2010) and schizophrenia (Eqbujo et al., 2016). Post-synaptically, many ID/NDD and SCZ risk genes regulate glutamate receptor signaling (Volk et al., 2015), as well as the dendritic spine actin cytoskeleton via effects on RHO GTPase and CAMKII signaling (Hall et al., 2015; Vissers et al., 2016). Thus, the synapse is a hot spot for neuropsychiatric disease risk gene pathology.

Here we investigated GIT1, a synaptic signaling and scaffolding protein (reviewed in Zhou et al., 2016). We propose that GIT1 plays central roles in synaptic neuropsychiatric disease risk gene pathology. Rare coding variants in *GIT1* are found in schizophrenia patients (Fromer et al., 2014; Purcell et al., 2014). Several of these schizophrenia-associated coding variants disrupt GIT1 activation of the p21-activated kinase PAK3 (Kim et al., 2016), a kinase with expression levels that are dysregulated in the brains of schizophrenia patients (Datta et al., 2015). Recently, a splice acceptor site variant in *GIT1* was found in a schizophrenia patient (Genovese et al., 2016). In addition, copy number variation at the *GIT1* locus, including both deletions and duplications, has been reported in patients with ID/NDD (Uddin et al., 2016). The fact that *GIT1* is a loss of function intolerant gene (Lek et al., 2016) strengthens the notion that these variants are pathological. Lastly, GIT1 has been reported to physically interact or form complexes with the schizophrenia risk gene products CNKSR2 (Lim et al., 2014; Ripke et al., 2014; CNSKR2 is also an ID risk gene—Houge et al., 2012), PTPRF (Ko et al., 2003; Ripke et al., 2014), and GRIA1 (Ko et al., 2003; Ripke et al., 2014), as well as the ID/NDD risk gene products ARHGEF6 (Kutsche et al., 2000; Bagrodia et al., 1999) and PAK3 (Allen et al., 1998; Bagrodia et al., 1999), all of which can be found at the synapse or at post-synaptic dendritic spines (Zhang et al., 2003; Collins et al., 2006; Zhang et al., 2005). These data strongly support the notion that GIT1 is a key protein that may regulate the function of multiple neuropsychiatric disease risk genes that may be involved in cognitive function.

GIT1 has been reported to function on both the pre-synaptic and post-synaptic sides of synapses (reviewed in Zhou et al., 2016). In pre-synaptic terminals, GIT1 binds to the active zone cytomatrix protein PCLO (Kim et al., 2003). Functionally, presynaptic GIT1 regulates the probability of vesicle release (Montesinos et al., 2015). On the post-synaptic side, GIT1 facilitates targeting of AMPA receptors to the post-synaptic density and the synaptic membrane (Ko et al., 2003). In addition, GIT1 plays key roles in dendritic spine formation and morphology (Zhang et al., 2003) by interacting with a number of proteins that regulate the actin cytoskeleton in spines, such as RAC1 and PAK3 (Zhang et al., 2005). With these multiple pre- and post-synaptic roles, GIT1 is a central regulator of synaptic transmission.

Given the key roles of GIT1 on both sides of synapses, involving interactions with ID/NDD and SCZ risk gene products, it is perhaps not surprising that *GIT1* loss of function in knockout mice causes cognitive deficits, such as decreased fear conditioning learning (Schmalzigaug et al., 2009), an absence of novel object recognition and reduced spatial learning and memory in the Morris water maze test (Won et al., 2011; Martyn et al., 2018), and impaired performance during operant learning (Menon et al., 2010). However, all previous studies in *GIT1* loss of function mice involved whole body knockout; these mice suffer from substantial post-natal lethality (Schmalziqaug et al., 2009) as well as impaired lung development (Pang et al., 2009). Thus, it is unclear whether cognitive deficits observed in whole-body GIT1 knockout mice are due to abnormal brain function versus general functional impairment due to pulmonary insufficiency.

Here we use conditional neural-specific *GIT1* knockout (NKO) mice to investigate cognitive deficits due to loss of function of *GIT1* in the nervous system. Unlike whole-body *GIT1* knockout, conditional neural-specific *GIT1* knockout does not cause post-natal lethality. We find that these mice have deficits in both fear conditioning learning and working memory. We then utilize a proteomic approach to analyze hippocampi from *GIT1*-NKO mice to identify protein phosphorylation events that depend on the presence of GIT1. We discuss the relevance of our findings for GIT1 regulation of protein networks involved in synaptic functions, and dysregulation of these networks and functions in neuropsychiatric diseases.

## Materials and Methods

### *GIT1* Conditional Neural Knockout Mice

Floxed *Git1* mice (Schmalzigaug et al., 2009; Jackson Laboratories Mouse Genome Informatics symbol Git1^tm1.1Rtp^) and Nestin-Cre (B6.Cg-Tg (Nes-cre)1Kln/J) mice were generously provided by Dr. Guoping Feng (MIT). Floxed *Git1* mice used here had been extensively (> 10 generations) backcrossed with C57BL/6 mice. Nestin-Cre mice were initially purchased from Jackson Laboratories (stock number 003771; genetic strain is the F2 hybrid of the closely related substrains C57BL/6 and C57BL/10) and then backcrossed with C57BL/6 mice. For breeding conditional *GIT1* neural-specific knockout mice according to the established protocol required to prevent germline transmission of the Nestin-Cre transgene (Bates et al.,1999), male *Git1*^flox/+^, *Cre^+^* transgenic mice were mated with female *Git1*^flox/flox^ *Cre^-^* mice to generate both neural knockout (GIT1-NKO; *Git1*^flox/flox^ *Cre^+^*) and control (*Git1*^flox/flox^ *Cre^-^*) males for behavioral testing. For some experiments (see below), Nestin Cre mice were purchased from Jackson Laboratories for use as additional controls in behavior experiments. Genotyping was carried out under standard conditions utilizing a previously published protocol (Schmalzigaug et al., 2009).

### Behavioral studies

Mice were group housed (4/cage) in a temperature and humidity controlled room and maintained on a reverse 12:12 light/dark cycle (7:00 am-7:00 pm) with *ad libitum* access to food and water. All behavior assays used male mice aged 8-12 weeks, and were performed during the light period under low level white light illumination. Experimental procedures were approved by the MIT institutional animal care and use committee under protocol 1012-102-15.

All behavioral studies were carried out with experimenter blinded to genotype. The last cage bedding change occurred ∼24 hours prior to the beginning of behavior testing. For all assays, mice were habituated in the test facility in their home cages for 1 hour prior to starting the task. Cohort 1 was tested in the following behavioral tests: T-maze, 3-chamber social interaction, elevated plus maze, pre-pulse inhibition, and fear conditioning with at least four days between each behavioral experiment. Follow up cohorts were run in T-maze and fear conditioning only as these demonstrated differences between control and *GIT1*-NKO mice in the initial cohort.

### T maze spontaneous alternation

Mice were placed in the start arm of the T-maze (5 cm wide x 28 cm long x 10 cm high; Stoelting) and allowed to freely explore the maze for 5 minutes. Arm entries were recorded and a correct alternation was defined as 3 consecutive arm entries into an arm not entered on the previous 2 trials. Chance performance in this task is 22%.

### Three-chamber social interaction

Male mice were used across all cohorts. Both stranger 1 and stranger 2 were wild type male S129 Sv males (Jackson laboratory) with matched age and body weight to test mice. Stranger mice were habituated by placing them inside an inverted wire cup for 30 minutes, two sessions per day for three consecutive days before experiments. Each stranger mouse was used maximally two times per day. The social test apparatus was made of a clear plexiglass box (65 (L) × 44 (W) × 30 (H) cm) with removable floor and partitions dividing the box into left, center, and right chambers. Center chamber (21 cm x 22 cm) is half the width of the left (21 cm x 44 cm) and right chamber (21 cm x 44 cm). These three chambers were interconnected with 5 cm openings between each chamber which can be closed or opened manually with a lever operated door. The inverted wire cups to contain the stranger mice were cylindrical, 10 cm in height, a bottom diameter of 10 cm with the metal bars spaced 0.8 cm apart. A weighted cup was placed on top of the inverted wire cups to prevent the test mice from climbing onto the wire cup. Each wire cup was used only one time per day then followed by extensive clean with 75% ethanol and water at the end of the test day. During the habituation phase, an empty wire cup was placed into left and right chamber, and the test mouse was placed into the middle chamber and allowed to explore for 15 minutes, with the doors into both side chambers open. During the sociability test phase, the test mouse was firstly gently introduced to the middle chamber with the doors to both side chambers closed, and an unfamiliar mouse (S1) was placed under the inverted wire cup in one of the side-chambers and a toy object (O) was placed under the inverted wire cup placed on the opposite side chamber. The location of the stranger mouse and object was counterbalanced between test trials to exclude side preference. The experimenter then lifted up the doorways to both side chambers simultaneously, and the test mouse was allowed to explore all three chambers for 15 minutes. During the social novelty test phase, the test mouse was again gently introduced into the middle chamber with the doors to both side chambers closed. Then a novel mouse (S2) was placed under the inverted wire cup, replacing the toy object (O) in one of the side-chambers. The experimenter then lifted up the doorways to both side chambers simultaneously, and the test mouse was allowed to explore all three chambers for an additional 15 minutes. Time spent in close proximity to the stranger mice or toy object was analyzed using Noldus Ethovison software.

### Elevated plus maze

Mice were placed in a closed arm of an elevated plus maze (Arm width = 10 cm, arm length = 50 cm, wall height = 30 cm, distance from floor = 55 cm; Coulbourn Instruments) and allowed to freely explore the maze for 5 minutes. Total time in the open and closed arms and the total distance travelled was recorded automatically via video tracking software (EthoVision).

### Acoustic startle response and prepulse inhibition

Each test chamber was equipped with a loudspeaker mounted 25 cm above the holding cylinder and a commercial startle reflex system (SR Lab, San Diego Instruments, CA). Individual mouse was placed inside the plexiglass holding cylinder mounted on a plexiglass platform. A piezoelectric accelerometer located beneath the platform was used to transform startle responses into units based on force and latency of startle. Data were collected at 250 samples/s and the maximum voltage attained on each trial was used as the dependent variable. Each test session started with a 5 minute acclimation period in the presence of 65 dB acoustic background noise followed by five 120 dB startle pulses. Pre-pulse trials followed the initial 120 dB startle acclimation. Each pre-pulse stimulation was 20 ms in duration, followed by a 40 ms startle stimulus of 120 dB. PPI was recorded for pre-pulse intensities of 70, 75, 80, 85 and 90 dB, and no stimulus. Each prepulse trial was administered ten times in a random order. Trials of 120 dB alone were randomly interspersed within the pre-pulse trials and used for comparison with the prepulse trials. The percent PPI was calculated using the formula [100−(response to pre-pulse + 120 dB)/(response for 120 dB alone)×100]. Acoustic startle trials were followed the PPI trials. Startle trials consisted of 40 ms pulses at 0 (no stimulus), 70, 75, 80, 85, 90, 95, 100, 105, 110, 115, and 120 dB. Each trial was presented five times in a randomized order.

### Fear Conditioning

Training: Mice were placed in one of two fear conditioning chambers (Coulbourn Instruments) and allowed to freely explore for 2 minutes (baseline). After 2 minutes, mice were presented with a 30s white noise (85dB) conditioned stimulus (CS) with a co-terminating 2s foot shock (0.75mA; US). Freezing was measured continuously for 2 minutes after the first CS-US presentation (immediate) which also served as the inter-trial interval (ITI). Mice were then presented with a second 30s CS co-terminating 2s US pairing, followed by 30 seconds before being removed from the training chamber. All freezing behavior was automatically scored by Freeze Frame software (Coulbourn) defined as a complete absence of movement other than respiration for a full second.

Context Test: 24 hours after initial training, mice were placed back into the original training chamber and freezing was measured continuously for 5 minutes (context).

Cued Test: One hour after contextual testing, the chambers were altered (grid floor covered with a smooth plastic floor (tactile); wall cues added (visual); vanilla extract scent added (olfactory)) and freezing was measured for 2 ½ minutes (Pre-CS). Then, the 85dB white noise was presented and freezing was measured continuously for 2 ½ minutes (cued).

### Behavioral Data Analysis

All behavioral data were collected by automated methods to remove potential confounds associated with human scoring, with the exception of spontaneous alternation in the T-maze, which was scored by the experimenter in real time, blinded to genotype. T-tests or repeated measures analysis of variance (ANOVA) were utilized where appropriate. All data were analyzed via SPSS v. 24

### Total and Phospho-proteomics

Total hippocampal proteins and phospho-proteins were detected by quantitative LC-MS/MS methods. Mass spectrometry detection and quantitation was performed in quadruplicate with paired samples consisting of male, 8-12 week old matched pairs (either littermates or age-matched) of control and GIT1-NKO mice. Following decapitation, mice heads were dipped in liquid nitrogen for 5 seconds to rapidly cool, but not freeze, the brain. Hippocampi were dissected rapidly on a cooling tray, and placed in Covaris bags (tissueTUBE TT05, Covaris), which were then flash frozen in liquid nitrogen. The frozen tissue was pulverized according to the manufacturer’s protocol on a Covaris CP02 Cryoprep Pulverizer. The pulverized tissue was lysed in 8M urea with protease and phosphatase inhibitors. The protein lysate (typical total yield ∼1-1.5 mg per hippocampus) was reduced, alkylated, and double digested with both Lys-C and Trypsin overnight. Equivalent amounts of tryptic peptides (∼1 mg) from each sample were labeled with TMT-10 reagent (Thermo Fisher Scientific) and the individual label incorporation was checked via LC-MS/MS. All samples had greater than 95% label incorporation. The labeled digests were combined and fractionated using basic reverse phase (bRP) chromatography (with an Agilent HPLC system) into 2 fractions for proteome analysis and 12 fractions for phosphopeptide analysis. Fractionation decreases sample complexity and increases the dynamic range of detection. From each fraction, 5% of the total volume was used for proteomic analysis while the remaining 95% was used for phosphopeptide enrichment using established protocol for Immobilized Metal Affinity Chromatography (IMAC) developed at the Broad Institute (Mertins et al., 2016; Mundt et al., 2018). The proteome and the phosphoproteome data were acquired on a Q-Exactive+ mass spectrometer (Thermo Fisher Scientific). Peptide spectrum matching and protein identification was performed using Spectrum Mill (Agilent). Peptide identification false discovery rates (FDRs) were calculated at 3 different levels: spectrum, distinct peptide, and distinct protein. Peptide FDRs were calculated in SM using essentially the same pseudo-reversal strategy evaluated by Elias and Gygi (Elias and Gygi 2010) and shown to perform the same as library concatenation. A false distinct protein ID occurs when all the distinct peptides that group together to constitute a distinct protein have a deltaForwardReverseScore ≤ 0. We adjusted settings to provide peptide FDR of 1-2% and protein FDR of 0-1%. SM also carries out sophisticated protein grouping using the methods previously described (Neshvizhskii and Aebersold 2005). Only proteins with >2 peptides and at least 2 TMT ratios in each replicate are counted as being identified and quantified. For each matched pair of control and *GIT1*-NKO hippocampi, Spectrum Mill calculated a fold-change for each protein or phospho-site (psite). Our initial criterion for a GIT1-regulated ‘hit’ was a requirement that fold-changes from all four matched sample pairs had to be coherent, i.e. in the same direction (up or down). We then normalized the mean of these four psite fold changes to the mean of the fold changes of the corresponding proteins induced by GIT1 neural knockout. For quantitating protein levels and fold changes, only peptides unique to individual proteins were counted; thus, peptides that are identical across multiple gene homologs were not counted. We then ranked our normalized psite fold change list by the magnitude of the change induced by GIT1 neural knockout. Lastly, we imposed a minimum fold change threshold of 25% (up or down); the 639 phospho-protein changes meeting these criteria are shown in **Supplemental Table 7**.

### GO category enrichment analyses

GO category enrichment analysis for all detected hippocampal proteins (**Figure 2B**) and GIT1-regulated phospho-proteins (**Figure 3B**) was performed using the Enrichment Analysis (PANTHER) web tool at www.geneontology.org, which output both fold-enrichment values and corrected false discovery rate p values (**Supplemental Table 2**). To generate the treemaps in **Figures 2B** and **3B**, we started with the indicated sets of enriched cellular component GO categories with corrected p values < 1e^-05^. We removed GO categories that could not be linked to a specific cellular component (e.g. “Cell” and “Intracellular”), and then we manually annotated each remaining GO category to its corresponding specific cellular component (e.g. the GO category “Chromatin” was annotated as being in the “Nucleus” component). Our manually annotated cellular components were the branches (shown as distinct colors on the treemap; branch names are shown as white text in each branch) of the treemap, with enriched GO categories as the leaves (individual lettered boxes in each branch). Leaf size was determined by the fold-enrichment value for each GO category. GO cellular component category enrichment among GIT1-regulated phospho-peptides (**Supplemental Figure 3; Supplemental Table 7**) was also quantitated and visualized with the BiNGO plugin (Maere et al., 2005) in Cytoscape version 3.5.1. The BiNGO visualization shows the GO cellular component categories as circular ‘nodes’ that were found to be statistically significantly enriched, superimposed on the corresponding regions of the GO hierarchy. Node size is proportional to the number mouse hippocampal phospho-peptides (**Supplemental File 1**) of GIT1-regulated phospho-protein (**Supplemental Figure 3**) genes that are annotated to that node’s GO cellular component category. Node color reflects the corrected p-value for enrichment: white nodes are not significantly enriched; colored nodes have significant p-values ranging from yellow (p between 0.001-1E^-7^) to dark orange (p < 1E^-7^).

**Figure 1.**
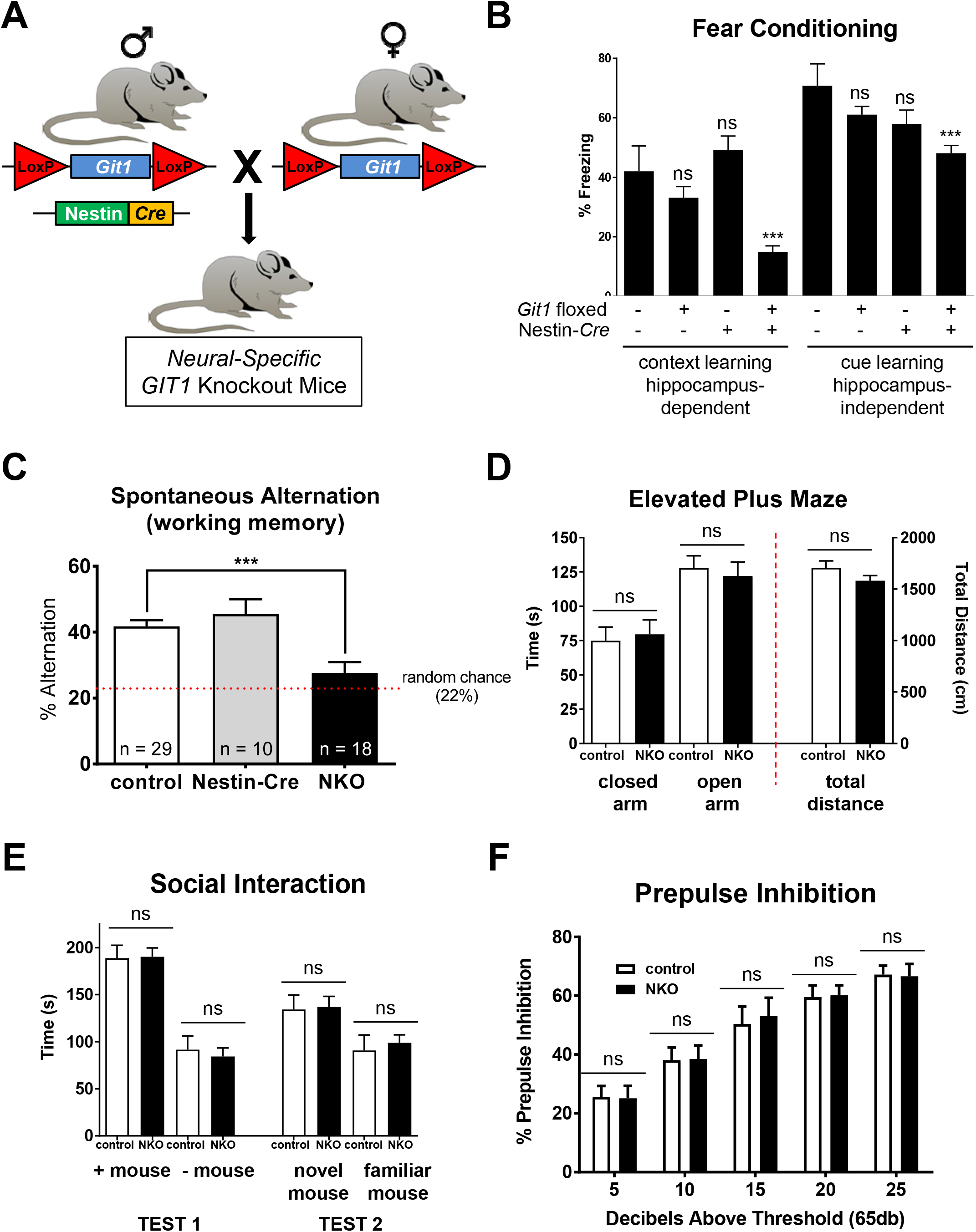
Cognitive deficits observed in *GIT1*-NKO mice tested in a battery of behavioral paradigms. **(A)** *GIT1*-NKO mice were bred by crossing males carrying the Nestin-Cre transgene that is expressed specifically in neural tissues, with females homozygous for floxed *GIT1* allele (see **Supplementary Figure 1**), resulting in neural-specific *GIT1* knockout. **(B)** Context and cued fear conditioning tests. N = 8 (GIT1^+/+^/Nestin-Cre^-^ control), 20 (GIT1^flox/flox^/Nestin-Cre^-^ control), and 18 (GIT1^flox/flox^/Nestin-Cre^+^ knockout). *** = p < 0.001; ns = not significant; one-way ANOVA followed by multiple comparison tests versus control. **(C)** T-maze spontaneous alternation test. Alternation was measured as the percentage of sets of three consecutive arm entries all occurring in non-previously visited arms over a 5 minute test period. *** = p < 0.001; ns = not significant; one-way ANOVA followed by multiple comparison tests versus control. **(D)** Elevated plus maze test. **(E)** Social interaction test. **(F)** Prepulse inhibition test.

**Figure 2.**
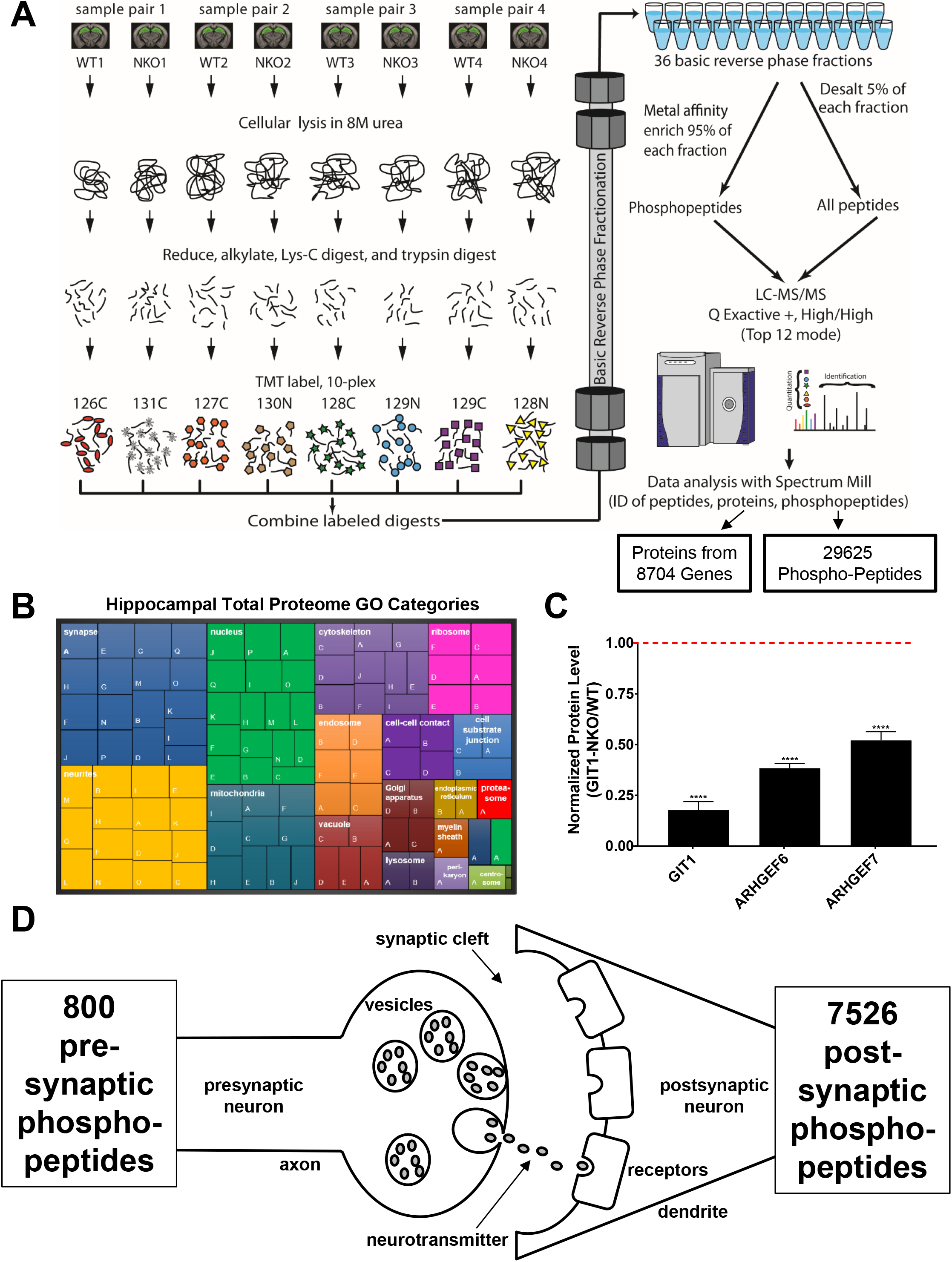
Phospho-proteomic analysis of GIT1-regulated synaptic phosphorylation signaling events within a network of neuropsychiatric risk genes. **(A)** Pipeline for discovery-based proteomic and phospho-proteomic profiling. Protein from dissected hippocampal (green region on coronal brain view) sample pairs was extracted with 8M urea and digested with LysC/trypsin. Peptides from each sample were labeled with chemical mass tag reagents for relative quantification; the 10-plex TMT labeling method is illustrated. Following labeling, peptides were combined and fractionated by basic reversed-phase (bRP) chromatography to decrease sample complexity and increase the dynamic range of detection. The global proteome of each plex was measured with 24 bRP fractions using 60 h (24x 2.5 h runs) of measurement time on a Thermo Q Exactive Plus instrument. For phospho-proteome analysis, phosphorylated peptides were enriched with immobilized metal affinity chromatography (IMAC) and injected as 12 LC-MS/MS runs requiring 30 h assay time per plex. **(B)** Treemap of enriched GO cellular component categories in the set of all proteins detected in the mouse hippocampus. Branches are shown as distinct colored areas with branch names in white text; leaves are lettered boxes within each branch, and specific leaf names are given as colored text in a few highlighted cases; for a complete list of leaf names see **Supplemental Table 2**. Leaf size corresponds to fold-enrichment values for each GO category. **(C)** Quantitated proteomic data indicating changes induced by *GIT1* neural knockout in the hippocampal levels of GIT1 and ARHGEF6/7. **** = p < 0.0001, unpaired t-test. Red dashed line indicates normalized level of protein in control hippocampi. **(D)**

**Figure 3.**
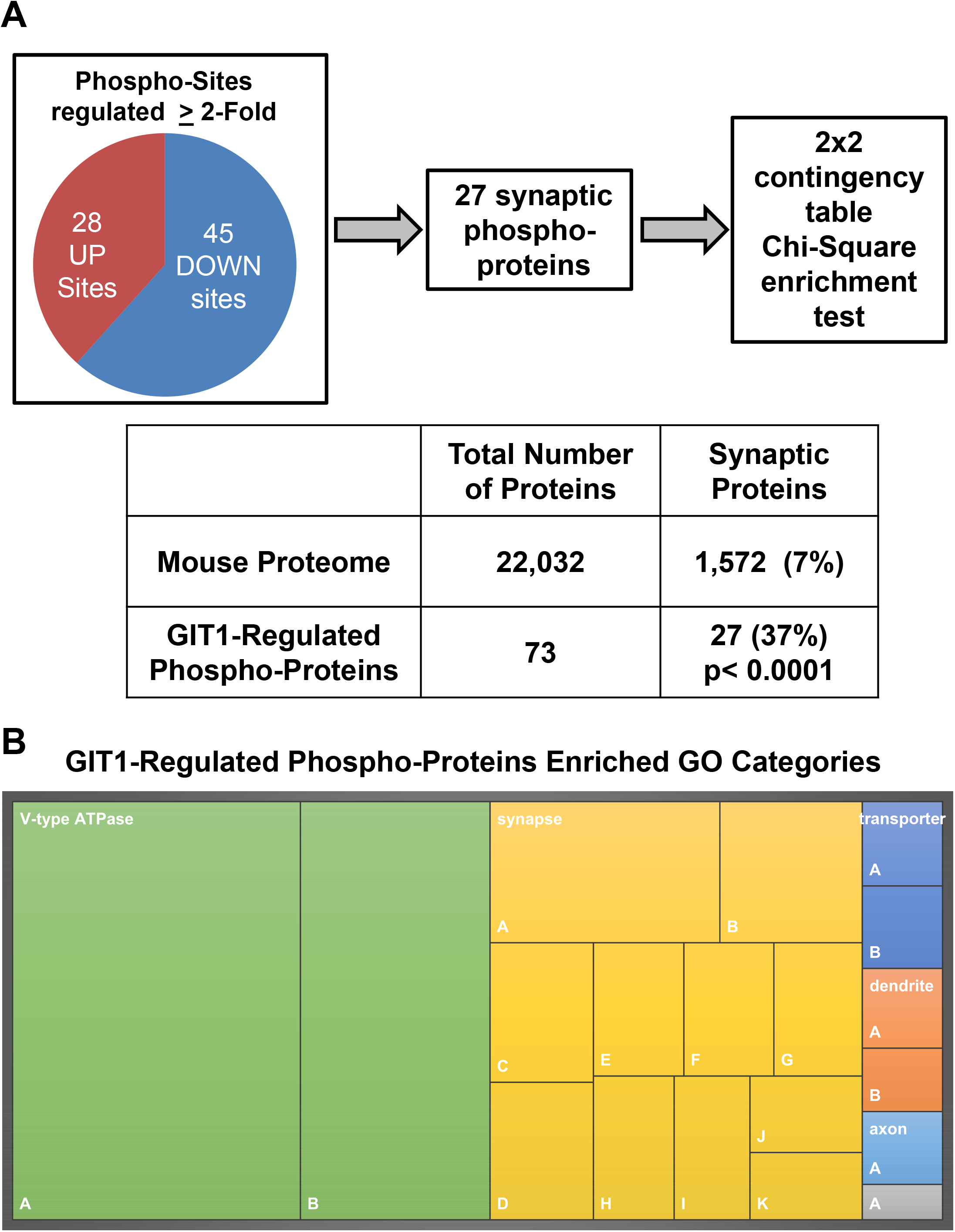
Global functional and synaptic analyses of mouse hippocampal proteome and phospho-proteome. **(A)** We cross-referenced our hippocampal phospho-peptide list with reported sets of biochemically purified pre-synaptic proteins (Boyken et al., 2013) and post-synaptic proteins (Collins et al., 2006) to estimate the potential number of distinct phospho-peptides on each side of the synapse. **(B)** Treemap of GO cellular component categories enriched in the set of all proteins detected in mouse hippocampus. Branches are shown as distinct colored areas with branch names in white text; leaves are lettered boxes within each branch, and specific leaf names are given as colored text in a few highlighted cases; for a complete list of leaf names see **Supplemental Table 10**. Leaf size corresponds to fold-enrichment values for each GO category.

### Molecular interaction and function enrichment database search

Molecular interaction and function (i.e. biological pathway) enrichment was performed using the expert curated biological pathway web tool Reactome (www.reactome.org). Using the ‘Analyze Data’ function, mouse protein sets (see Results) were entered, converted to human equivalents, and evaluated for pathway over-representation using default settings.

## Results

### Cognitive deficits in neural-specific conditional *GIT1* knockout mice

To analyze the behavioral consequences of GIT1 dysfunction in the context of intact neural circuits *in vivo*, we utilized a *GIT1* knockout mouse. Three distinct alleles generating whole-body *GIT1* knockout mice have been created, and mice with these knockout alleles have several learning and memory deficits (Schmalzigaug et al, 2009; Menon et al., 2010; Won et al., 2011). However, whole-body *GIT1* knockout mice also have greatly reduced postnatal survival (Schmalzigaug et al., 2009; Won et al., 2011) and impaired cardiac, pulmonary, and skeletal dysfunction (Pang et al., 2009; Menon et al., 2010; Pang et al., 2011) potentially confounding the interpretation of behavioral data in these mice. To overcome these previous limitations and create a model system to investigate the neural-specific functions of GIT1, we utilized the Cre/lox system by crossing Nestin-Cre expressing driver mice with one of the previously generated mouse strains carrying floxed *GIT1* alleles (Schmalzigaug et al., 2009) to conditionally inactivate *GIT1* in the central and peripheral nervous systems (*GIT1*-NKO mice; **Figure 1A**; **Supplemental Figure 1**). Analysis of adult hippocampal tissue from these neural-specific *GIT1*-NKO mice showed almost undetectable expression of GIT1 (**Supplemental Figure 1**). These mice had normal post-natal survival rates in contrast to the previously reported whole-body *GIT1* knockout mice, but were slightly smaller in overall body size similar to the whole-body *GIT1* knockout mice (data not shown).

To broadly assess the behavioral consequences of neural-specific loss of GIT1, *GIT1*-NKO mice were put through a battery of tests. To assess cognition, *GIT1*-NKO mice were first tested in both contextual fear conditioning, which assesses a memory process dependent on both the hippocampus and the amygdala (Kim & Fanselow, 1992; Muller et al., 1997), and cued fear conditioning, which assesses amygdala-dependent memory processes (Muller et al., 1997). **Figure 1B** shows that *GIT1*-NKO mice exhibit both hippocampus- and amygdala-dependent fear conditioning deficits when compared to Nestin-Cre expressing, non-floxed *GIT1*^+/+^ control mice. This did not simply reflect a difference in locomotor activity levels, as *GIT1*-NKO mice had no difference in freezing responses during the training period as compared to control mice (p = 0.1315; two-tailed t-test). Next, the mice were tested in a spontaneous alternation cortex- and hippocampus-dependent working memory T-maze task (Lalonde, 2002; Dudchenko, 2004). **Figure 1C** shows that *GIT1*-NKO mice had virtually no ability to recall arm visits in the T-maze, suggesting a severe working memory deficit. In contrast, *GIT1*-NKO mice had no deficits in social interaction (**Figure 1E**, three-chamber social interaction test, reviewed in Silverman et al., 2010) or prepulse inhibition (**Figure 1F**), a test of sensorimotor gating, which is defective in schizophrenia patients (e.g. Braff et al., 1978). In addition, *GIT1*-NKO mice scored equivalent to control mice in the elevated plus maze test (**Figure 1D**), indicating that these mice have normal levels of anxiety. These results demonstrate that GIT1 function is critical specifically for cognitive learning and memory processes, and are in accordance with deficits in contextual and cued fear conditioning reported previously for whole-body *GIT1* knockout mice.

### Proteome-wide identification of GIT1-regulated phosphorylation events in hippocampus

GIT1 regulation of synaptic processes might occur simply by scaffolding to facilitate the proximal localization of key synaptic machinery components required for function. However, given the ability of GIT1 to regulate multiple kinase signaling cascades (Zhou et al., 2016), GIT1 may also modulate synaptic processes by controlling the phosphorylation states of key synaptic proteins. To address this question in an unbiased manner, we employed mass spectrometry-based proteomics to discover GIT1-regulated phosphorylation events in mouse hippocampus. Notably, this analysis was proteome-wide—i.e. not restricted to synaptic proteins—as we used whole hippocampal lysate as the input into the proteomic pipeline. To preserve protein phosphorylation states in the hippocampi, mouse heads were quickly chilled after decapitation by a brief dip in liquid nitrogen (Jordi et al., 2013). Following dissection on a surface kept at −20 degrees, hippocampi were flash frozen in liquid nitrogen and then put through a standardized lysis process optimized for phospho-proteomics (Mertins et al., 2016; Mundt et al., 2018; and see methods). Hippocampal lysates from wild-type and *GIT1*-NKO mice (four biological replicates of each) were then subjected to immobilized metal affinity chromatography enrichment-based phospho-peptide mass spectrometry detection and quantitation, as well as parallel proteomic detection and quantitation of total protein levels (**Figure 2A**).

To our knowledge, these proteomic data represent the most comprehensive survey to date of proteins and phospho-proteins expressed in the hippocampus (**Figure 2A**). We found that 8704 genes expressed at least one characterized protein product; these genes comprise nearly 40% of the protein coding genes in the mouse genome according to Gencode (http://www.gencodegenes.org/mouse_stats.html) (**Supplemental Table 1**). Confirming an expected expression pattern, these proteins include receptors for the major neurotransmitter systems that have been described in the hippocampus: glutamate, GABA, acetylcholine, dopamine, serotonin, and norepinephrine (**Supplemental Table 1**; Vizi & Kiss, 1998; González-Burgos & Feria-Velasco, 2008). Gene ontology (GO) cellular component category enrichment analysis (using PANTHER; **Figure 2B and Supplemental Table 2**) showed the expected over-representation of proteins involved in synapses and neurites, and also highlighted the important roles of the nucleus, mitochondria, and the cytoskeleton, as well as other cellular components, in cells of the hippocampus. Similar GO category enrichment analyses for biological process and molecular functions are shown in **Supplemental Table 2**. Interestingly, our data support the notion of a role for the hippocampus in schizophrenia pathology (Harrison, 2004); of the 58 single-gene GWAS loci containing variants associated with schizophrenia risk identified by Ripke et al (2014), 36 of these express proteins we detect in the hippocampus (**Supplemental Table 6**). Lastly, as a key control, our proteomic data showed the expected large decrease in GIT1 levels in the neural knockout mice’ hippocampi (**Figure 2C**; remaining residual levels of GIT1 are likely from vasculature). In addition, the GIT1-associated PIX proteins (ARHGEF6 and ARHGEF7) are downregulated by GIT1-NKO; this lack of stability of PIX proteins in the absence of GITs has been observed previously (Won et al., 2011; Zhou et al., 2016). These data underscore the variety and complexity of protein expression and function in the hippocampus.

While it is well-appreciated that regulation of protein phosphorylation is a key aspect of synaptic plasticity (Woolfrey & Dell’Acqua, 2015), our data also indicate the staggering complexity of protein phosphorylation in the hippocampus, with 29,625 unique phospho-peptides detected on 6596 proteins, many of which had multiple phosphorylation sites/states (**Supplemental Table 3**). By cross-referencing these phospho-peptides with published lists of synaptic proteins (Boyken et al., 2013; Collins et al., 2006), we can deduce that, at synapses, there may be up to 800 and 7526 distinct phospho-peptides (many synaptic proteins contain multiple phospho-peptides) on the pre- and post-synaptic sides, respectively (**Figure 2D**). Searching for these phosphorylation states in the PhosphoSite public protein post-translational modification database (www.phosphosite.org) revealed both known and previously unreported phosphorylation events, reflective of the depth of the proteomic analysis performed. For example, 14 phosphorylation sites were detected in GIT1, and all of those are present in the PhosphoSite database (**Supplemental Table 4**). In contrast, 35 phosphorylation sites were detected in the schizophrenia risk gene product GRIN2A, and 7 of those were apparently novel (i.e. not present in the PhosphoSite database; **Supplemental Table 4**). Global analysis of Gene Ontology (GO) cellular component category enrichment in the total set of all phospho-proteins in our data (**Supplemental Table 3**) using the BiNGO app in Cytoscape (http://www.cytoscape.org/) revealed 275 enriched (p < 0.01) GO categories, comprising many basic and neuron-specific cellular components (**Supplemental Table 5;** hierarchical network visualization in **Supplemental File 1**).

Parallel proteomic detection and quantitation of total protein levels allowed for normalization of phospho-peptide levels to the corresponding levels of total protein in wild-type and *GIT1*-NKO samples; these normalized phospho-peptide levels were then used to quantitate phosphorylation state changes induced by *GIT1* knockout. Our initial criteria for a ‘hit’ in this assay were: 1) a *GIT1* knockout-induced change in phosphorylation state in the same direction (up or down) across all four of our paired replicate samples; and 2) a minimum change up or down of 25%. A total of 639 protein phosphorylation state changes, of at least 25% up or down, were observed consistently across all samples (**Supplemental Table 7**; see Methods section for details of quantitation method).

Some of the proteins with phosphorylation states altered by *GIT1* knockout are known to directly interact or associate with GIT1, including ARHGEF6 (Zhou et al., 2016), GIT2 (Kim et al., 2003), PCLO (Kim et al., 2003), CNKSR2 (Lim et al., 2014), CAMK2A and CAMK2B (Pang et al., 2008), PTK2 (Zhao et al., 2000), MYO18A (Hsu et al., 2010), PPFIA2 (Ko et al., 2003), RIMS1 (Shin et al., 2003), NCK1 (Frese et al., 2006), and ERC2 (Ko et al., 2003b); or to associate with ARHGEF6/7, including PAK1 and PAK2 (Bagrodia et al., 1999), SHANK1 (Park et al., 2003), and NSF (Martin et al., 2006). Indeed, we have replicated several of these reported interactions with GIT1 (**Supplemental Figure 2**). The GIT1 interactor NCK1 also interacts with the kinase TNIK (Fu et al., 1999), and TNIK phosphorylates serine 315 in CAMK2B (Wang et al., 2016); serine 315 phosphorylation was upregulated by *GIT1* knockout, suggesting that the GIT1-NCK1 interaction may inhibit TNIK. Supporting this notion, phosphorylation of a δ-catenin serine targeted by TNIK (serine 528 in CTNND2; Wang et al., 2016) was also slightly upregulated by *GIT1* knockout (by ∼16%, which was less than the 25% threshold we set for ‘hits’; data not shown). Taken together, these data indicate that phosphorylation states of proteins may be regulated via their interactions with the GIT1-ARHGEF6/7 complex and kinases that are regulated by this complex.

Because GIT1 stimulates the activity of p21-activated kinases (PAKs) and microtubule-associated kinase (MAPK) (Kim et al., 2017), and GIT1 has been reported to scaffold MEK-ERK MAP kinases (Yin et al, 2004; Yin et al, 2005; Zhang et al, 2009), we anticipated that *GIT1* knockout would decrease the phosphorylation of known PAK and/or MAPK targets. Indeed, the phospho-site that was the most downregulated by *GIT1* knockout, serine 92 on SEPT3, may be a PAK target, as PAKs phosphorylate septins in yeast (Versele & Thorner, 2004). In addition, another one of the strongest effects of *GIT1* knockout was a decrease in phosphorylation of the microtubule destabilizer STMN1 at serine 63; this serine is known to be a target of PAK (Wittmann et al., 2004). Likewise, *GIT1* knockout also decreases the phosphorylation of ARHGEF2, ARHGDIA, and SPEN, which are known targets of PAK (Aslan et al., 2013; Shin et al., 2009, Vadlamudi et al., 2005). Phosphorylation of PAK1 serine 144 and PAK2 serine 141, which get autophosphorylated upon PAK1/2 binding to GIT1/PIX (e.g. Kim et al., 2017), were downregulated by *GIT1* knockout. MYO18A phosphorylation was also decreased by *GIT1* knockout; MYO18A is a known PAK interactor (Hsu et al., 2010). Lastly, several other sites of phosphorylation decreased by GIT1 knockout (such as NSF serine 207, NSMF serine 72, DIRAS2 serine 35, and SCN1A serine 607) match, or nearly match, key characteristics of optimal PAK phospho-acceptor sites (R at position −2 and large hydrophobic residues—W, I, V, Y—at +1 to +2 relative to the phosphorylated serine; Rennefahrt et al., 2007; these phospho-sites are highlighted in **Supplemental Table 7**). In addition, the phosphorylation of numerous KSP motifs in neurofilament H was decreased by *GIT1* knockout; these motifs can be phosphorylated by MAPK (Veeranna et al., 1998). Thus, known GIT1 interactions and/or PAK/MAPK targets account for a subset of the GIT1-regulated phosphorylation state changes observed here; for the remainder, mechanisms of regulation are unknown, and these may represent previously undiscovered targets of PAKs, MAPKs, or other kinases.

### Biological network enrichment analysis of GIT1-regulated phospho-proteins

We detected 73 proteins with phosphorylation states that were robustly regulated (more than 2-fold) by *GIT1* knockout: 45 were downregulated, and 28 were upregulated (**Figure 3A**). We tested whether synaptic proteins were enriched in this set of GIT1-regulated phosphoproteins. We used the union of lists of biochemically purified pre-synaptic proteins (Boyken et al., 2013) and post-synaptic proteins (Collins et al., 2006) as a reference list of ‘synaptic proteins’, which totaled 1572 proteins, or ∼7%, out of a total of 22032 protein-encoding genes in the mouse genome according to Gencode (http://www.gencodegenes.org/mouse_stats.html). The occurrence of synaptic phospho-proteins in our GIT1-regulated set by pure random chance would therefore be 5 (i.e. 7% of 73). In fact, we observed 27 synaptic proteins among the set of phospho-proteins regulated at least 2-fold by GIT1. This enrichment of synaptic phospho-proteins regulated by GIT1 was statistically significant (**Figure 3A**; p < 0.0001 by Chi-square test with Yates correction). Accordingly, GO cellular component enrichment analysis (using PANTHER) among GIT1-regulated hippocampal phospho-proteins (**Figure 3B**) showed that the vast majority of the enriched categories, as ranked by the degree of fold enrichment, involved V-type ATPases and synaptic components (see **Supplemental Table 9** for a list of all enriched cellular component categories; the cell compartment specific subset of these categories visualized the in treemap in **Figure 3B** is listed in **Supplemental Table 10**). Indeed, the V-type ATPase categories may also be considered as synaptic components, as V-type ATPases function to facilitate loading of synaptic vesicles with neurotransmitters (Edwards, 2007). These data indicate that one of GIT1’s primary roles in the hippocampus is to regulate levels of synaptic protein phosphorylation.

Inspection of the set of synaptic proteins with phosphorylation states regulated by GIT1 indicated that these proteins fell into the same functional categories as has been reported for GIT1-associated proteins in the brain (Zhou et al., 2016). Of synaptic phosphoproteins changed by more than 2-fold, 9 are involved in presynaptic vesicle functions, 10 are involved in post-synaptic density functions, and 16 regulate the cytoskeleton. For example, phospho-SEPT3 (serine 92) was downregulated by more than 70% by *GIT1* knockout. SEPT3 is known to be a phosphoprotein localized to presynaptic nerve terminals, and it appears to be involved in synaptic vesicle recycling (Xue et al., 2004). Phosphorylation of serine 499 of the post-synaptic density scaffolding protein SHANK1 was upregulated more than 3.1-fold. DOCK10 phosphorylation at tyrosine 1980 was downregulated by nearly 58%. DOCK10 is a dendritic spine cytoskeleton regulator that acts through effects on CDC42, N-WASP, PAK3, and RAC1 (Jaudon et al., 2015). Phosphorylation of another neuronal cytoskeleton regulator, MACF1 (serine 7245), was downregulated by 56%. Both DOCK10 and MACF1 have been reported to regulate dendrite (or dendritic spine) and axon growth (Jaudon et al., 2015; Ka & Kim, 2016). Thus, GIT1 regulates the phosphorylation states of synaptic proteins associated with the functions of synaptic vesicles, the post-synaptic density, the neuronal cytoskeleton, and axon and dendrite growth.

To identify the protein networks with phosphorylation states dependent on the presence of GIT1 in an unbiased manner, we performed pathway enrichment analysis with the Reactome curated pathway database (www.reactome.org). This tool uses a set of user input seed proteins, in the present case identified in our unbiased proteomics experiment, to search a manually curated database covering protein-protein interactions across a wide swath of human molecular physiology and disease, and then generates error-corrected false discovery rate (FDR) values for pathway over-representation. Included in this database are brain pathways that are specific to neurons in general, specific types of neurons (defined by neurotransmitter type), and glia. Initially, we tested a seed set consisting of all phospho-proteins identified in control or GIT1-NKO hippocampi (6596 phospho-proteins, many of which were phosphorylated at multiple sites; see **Supplemental Table 3**). No pathways were enriched in this ‘total hippocampal phospho-proteome’ seed set; all pathways had a FDR value > ∼0.9. We next used a seed set consisting of all phospho-proteins regulated by more than 30% (up or down; 512 phospho-proteins; see **Supplemental Table 7**) by *GIT1* neural knockout. With this ‘GIT1-regulated hippocampal phospho-proteome’ seed set, four pathways had FDR values < 0.05 (**Supplemental Table 8**). The most statistically enriched pathway was ‘Neuronal system’, consistent with the notion that GIT1 regulates phospho-proteins in neurons, and indicating that the Reactome database is sufficiently annotated to detect organ-specific pathways. The other statistically significantly enriched pathways involved molecular interactions that connect and support synapses (e.g. neurexin-neuroligin interactions). A number of nominally significant pathways (FDR < 0.1) involved mechanisms of neurotransmission at synapses common to several neurotransmitter types, in particular including excitatory glutamatergic synapses. A growing body of genetic research implicates both neurexin-neuroligin interactions and excitatory glutamatergic synapses with cognition and neuropsychiatric disease (Doherty et al., 2012; Hall et al., 2015). Thus, these unbiased Reactome-generated observations of statistically enriched pathways involving phospho-proteins regulated by GIT1 suggest that defects in GIT1 signal transduction functions at excitatory synapses may underlie pathology in neuropsychiatric diseases.

### Schizophrenia risk genes in GIT1-regulated synaptic phospho-protein networks

Having identified GIT1-regulated phospho-proteins at the synapse, we next asked whether any of these proteins have been associated with risk for schizophrenia. To address this question, we intersected the list of GIT1-regulated synaptic phospho-proteins (**Supplemental Table 7**) with schizophrenia risk genes identified in a study of common variation in schizophrenia patients (Ripke et al., 2014). This analysis revealed three GIT1-regulated synaptic phospho-proteins that have common variants that are associated with risk for schizophrenia: RIMS1, GRIN2A, and CNKSR2. Notably, GIT1 has been reported to directly interact with one of these proteins: CNKSR2 (Lim et al., 2014); we have been able to reproduce observation of this interaction using a co-immunoprecipitation assay in HEK293 cells (**Supplemental Figure 2**). The effect of GIT1 knockout on the phosphorylation status of these three proteins is shown in **Figure 4A** (black bars). Notably, two of these GIT1-regulated, schizophrenia-associated proteins (CNKSR2 and GRIN2A) are also associated with intellectual disability (Endele et al., 2010; Houge et al., 2012). We next constructed pre- and post-synaptic GIT1-schizophrenia risk gene interaction networks (**Figure 4B and C**) using GIT1 and RIMS1 (presynaptic), or GIT1 and CNKSR2 (postsynaptic) as starting nodes, and then adding interacting nodes from the sets of GIT1-regulated phospho-proteins (**Supplemental Table 7**) known to be expressed in those two synaptic compartments, particularly in excitatory glutamatergic synapses (Boyken et al., 2013; Collins et al., 2006), with interactions (edges) from the BioGRID protein interaction database (https://thebiogrid.org/). The effect of *GIT1* knockout on phosphorylation of a subset of these pre- and post-synaptic proteins is shown in **Figure 4A** (grey bars). Several proteins in these networks are known to be encoded by autism (Sanders et al., 2015) or developmental delay (McRae et al., 2017) risk genes (marked in **Figure 4B**) and it is possible that several more of the genes encoding these networks’ proteins will be identified as schizophrenia risk genes in future studies as schizophrenia GWAS and sequencing sample sizes increase. Future work will be required to determine how these phosphorylation changes affect the functioning of GIT1-regulated pre- and post-synaptic networks, and the behavioral consequences of the effects on these networks; however, the current literature contains some clues. For example, GIT1 knockout dramatically induces phosphorylation of CAMK2B at threonine 325, which is a residue in its actin-binding domain; this may allow CAMK2B to dissociate from actin during signaling events that induce synaptic plasticity and dendritic spine actin remodeling (Kim et al., 2015), suggesting that GIT1 may function to negatively regulate synaptic plasticity-associated dendritic remodeling through suppression of CAMK2B phosphorylation. In addition, mutation of several phosphorylated residues in GRIN2A to alanine, including GIT1-regulated S1291 and Y1292, caused deficits in Y-maze working memory (Balu et al., 2016) similar to the effect of GIT1 neural knockout observed here (**Figure 1C**). The identification of pre- and post-synaptic networks containing GIT1-regulated phospho-proteins, including schizophrenia and intellectual disability risk genes, clearly suggests the central role of GIT1-regulated proteins in a hot spot for neuropsychiatric disease pathology—the excitatory glutamatergic synapse.

**Figure 4.**
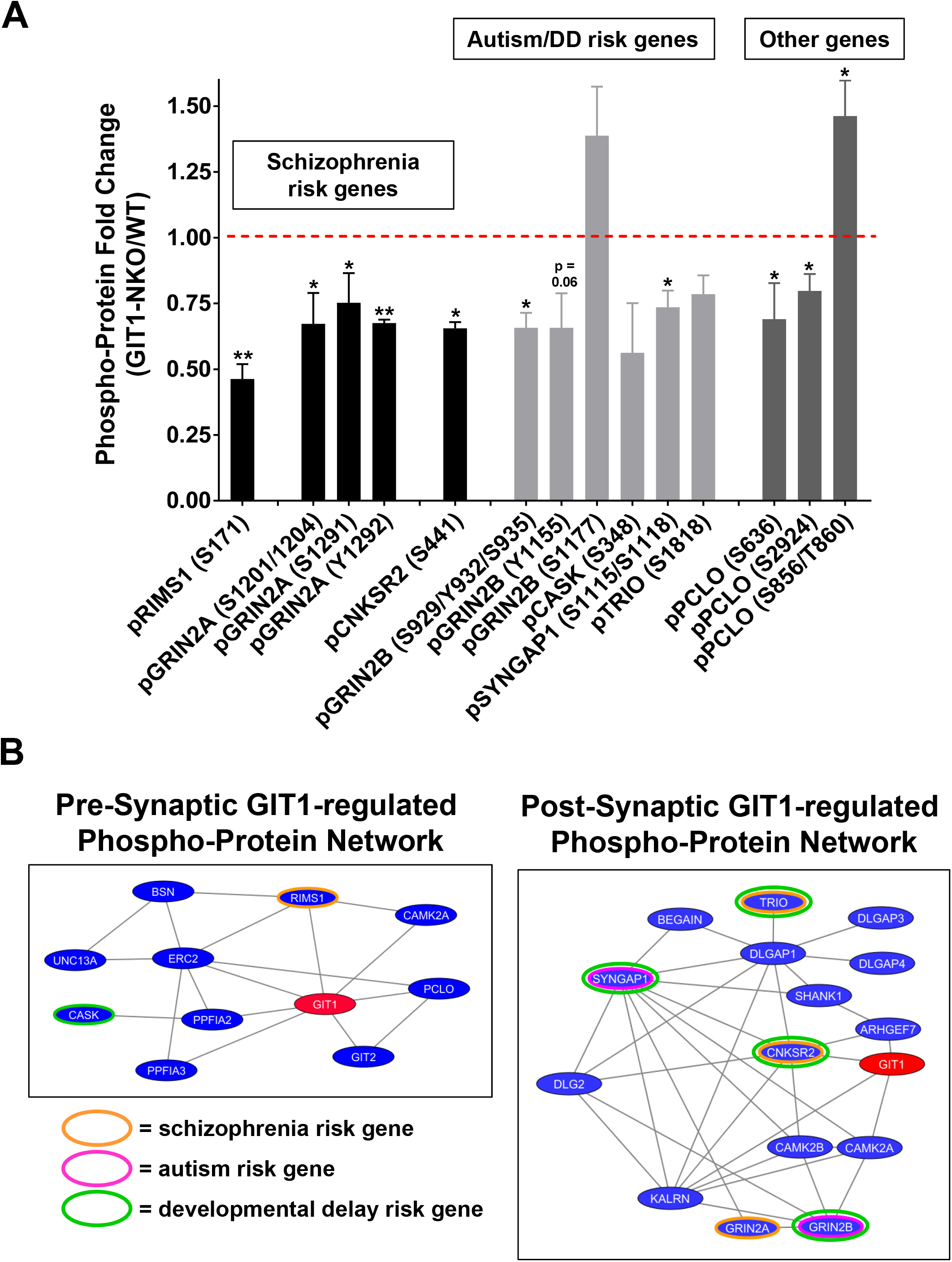
Enrichment of synaptic proteins in the set of hippocampal phospho-proteins regulated by GIT1. **(A)** Hierarchical organization of over-represented GO cellular component categories among GIT1-regulated hippocampal phospho-proteins. Statistical over-representation calculation and hierarchy mapping were performed using the BiNGO tool in Cytoscape software. Red circles indicate specific GO categories discussed in the text (see Results). **(B)** Statistical 2 × 2 contingency table Chi-square test of enrichment of synaptic proteins in the set of phospho-proteins robustly regulated (> 2-fold) by *GIT1* neural knockout. Synaptic proteins were defined as the union of the presynaptic and postsynaptic proteins identified in Boyken et al., 2013 and https://www.genes2cognition.org/.

**Figure 5.** GIT1-regulated synaptic networks affected in schizophrenia. **(A)** Effect of *GIT1* knockout on phosphorylation of four schizophrenia risk gene proteins. * = p < 0.05; ** = p < 0.01; p values output by Spectrum Mill for the indicated phospho-protein sites. **(B)** Model of the synaptic functions of GIT1-regulated schizophrenia risk gene proteins.

## Discussion

Building on recent efforts to elucidate the genetic architecture of neuropsychiatric diseases, the delineation of components, interactions, and signal transduction flow in key molecular networks and pathways that are disrupted in these diseases is a critical next step for the field. Here we study one such component with function and expression altering genetic variation in schizophrenia and neurodevelopmental disorders: the synaptic scaffolding and signaling protein GIT1. We utilize behavioral testing and phospho-proteomic analysis in mice with conditional neural-specific knockout of *GIT1* to identify aspects of cognition, as well as hippocampal phosphorylation signaling events that are regulated by GIT1. We find that *GIT1* neural knockout mice have cognitive deficits in associative learning and working memory, while social interaction, sensorimotor gating, and anxiety behaviors were normal. Our proteomic analyses indicate that GIT1 regulates phosphorylation states within networks of pre- and post-synaptic proteins. Excitingly, this includes a set of proteins within these synaptic networks that have been implicated as risk factors in genetic studies of schizophrenia and neurodevelopmental disorders. These data suggest that deficits in GIT1 functioning in synaptic networks may contribute to cognitive dysfunction in neuropsychiatric diseases.

Copy number variation at the *GIT1* locus, including both deletions and duplications, has been reported in patients with neurodevelopmental disorders (Uddin et al., 2016); these patients generally have developmental delay and intellectual disability. Here we identified cognitive deficits in a mouse model with complete nervous system-specific knockout of *GIT1*. Previous studies reporting associative learning deficits in *GIT1* knockout mice (Schmalzigaug et al., 2009; Won et al., 2011; Menon et al., 2010; Martyn et al., 2018) utilized whole-body knockout, which produces post-natal cardiovascular dysfunction and lethality (Pang et al., 2009; Schmalzigaug et al., 2009). In contrast, mice studied here with neural-specific *GIT1* knockout had normal post-natal survival. Consistent with whole-body knockout mice, neural-specific *GIT1* knockouts had an associative learning deficit in the fear conditioning test (**Figure 1**). In addition, we found that neural-specific *GIT1* knockout mice had a severe working memory deficit in the spontaneous alternation test (**Figure 1**). These findings suggest functional deficits in both the hippocampus and cortex, which are key brain regions involved in associative learning and working memory (Kim & Fanselow, 1992; Lalonde, 2002).

To begin to elucidate the molecular mechanism of GIT1’s role in learning and memory, we performed phospho-proteomic analysis of GIT1-regulated signaling in the hippocampus. The set of hippocampal phospho-proteins regulated by GIT1 contained many synaptic proteins that fell within several functional categories: presynaptic vesicle trafficking, post-synaptic receptor clustering, and post-synaptic dendritic spine cytoskeleton regulation. Excitingly, several GIT1-regulated hippocampal phospho-proteins in these functional categories have common genetic variants associated with schizophrenia: RIMS1, GRIN2A, and CNKSR2. (The complete list of GIT1-regulated phospho-proteins that are schizophrenia GWAS genes is shown in Supplemental Table 11.) These data link GIT1 to an apparent hotspot for schizophrenia pathology—the excitatory glutamatergic synapse.

GIT1 regulation of phosphorylation of the presynaptic vesicle trafficking protein PCLO is especially intriguing in light of the recently reported role of GIT1 in negatively regulating presynaptic vesicle release probability, possibly by regulating actin dynamics at the active zone (Montesinos et al., 2015). PCLO also appears to negatively regulate synaptic vesicle exocytosis via effects on actin (Waites et al., 2011). We hypothesize that GIT1-regulated phosphorylation of PCLO may facilitate PCLO’s actions on the active zone actin cytoskeleton. This effect may be modulated by neuronal activity, as depolarization also regulates a subset of the GIT1-regulated phospho-sites in PCLO (Kohansal-Nodehi et al., 2016). These will be important questions to address in future studies.

GIT1 regulation of phosphorylation of the schizophrenia-associated NMDA receptor subunit GRIN2A may play a role in the working memory deficit we observed in the spontaneous alternation test. Two GRIN2A phosphorylation sites that had decreased levels of phosphorylation in *GIT1*-NKO mice (serine 1291 and tyrosine 1292) may be required for performance of this task—Balu and colleagues (2016) found that mice with these two residues mutated to alanine and phenylalanine, respectively, showed a deficit in the spontaneous alternation test.

In schizophrenia patients, several rare coding/splicing variants of *GIT1* have been identified (Fromer et al., 2014; Purcell et al., 2014; Genovese et al., 2016). Some of these variants impair GIT1 signaling via activation of PAK3 kinase (Kim et al., 2016). To date, common variation at the *GIT1* locus has not been associated with risk for schizophrenia (Ripke et al., 2014); however, as shown here, GIT1-regulated molecular networks include a number of genes at loci at which common variation has been associated with schizophrenia risk. To assess the effects of rare coding variants on GIT1-associated molecular networks, it will be necessary to study model systems expressing these variants, such as transgenic mice, or human induced pluripotent stem cell-derived neurons from patients with these coding variants. These studies are currently under way in our laboratory.

## Acknowledgments

We thank members of the Stanley Center for Psychiatric Research and the Chemical Neurobiology Laboratory for helpful discussions and critical feedback. Dr. Guoping Feng is thanked for generously sharing GIT1 mice and for his critical feedback. This work was supported by funding from the Stanley Medical Research Institute, the National Institute of Mental Health (R01MH095088), and the Stuart & Suzanne Steele MGH Research Scholar Award (SJH).

---

**Supplemental Figure 1:**
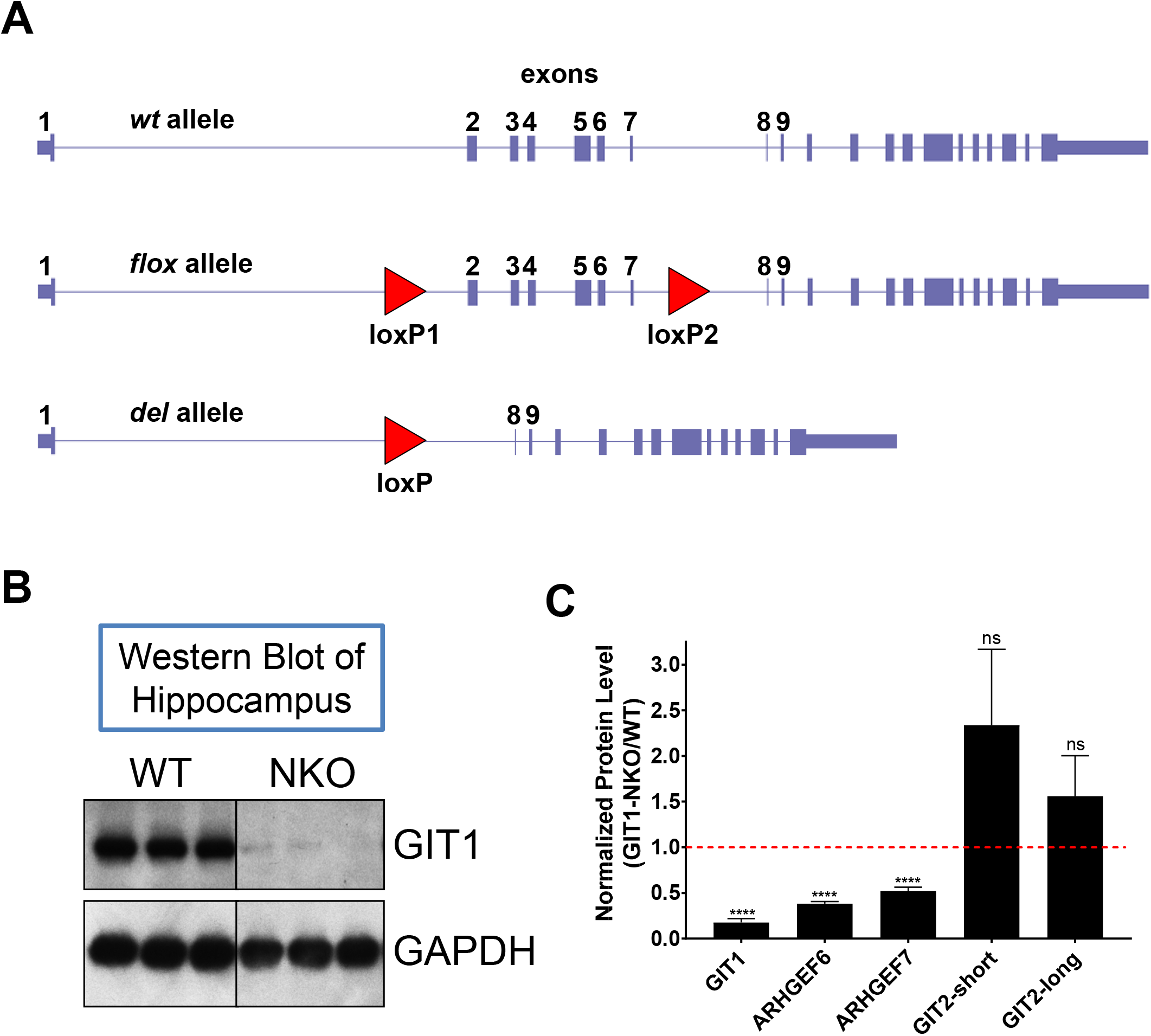
Diagram of *GIT1* conditional neural KO mouse and western blot showing loss of GIT1 expression from brain regions. GIT2 (both long and short isoforms) was observed to be upregulated potentially as compensation for the loss of GIT1 protein. However, this apparent compensatory upregulation of GIT2 was unable to replace GIT1’s role in cognitive behaviors (Figure 1), supporting the contention that GIT1 and GIT2 have different roles in the brain.

**Supplemental Figure 2:**
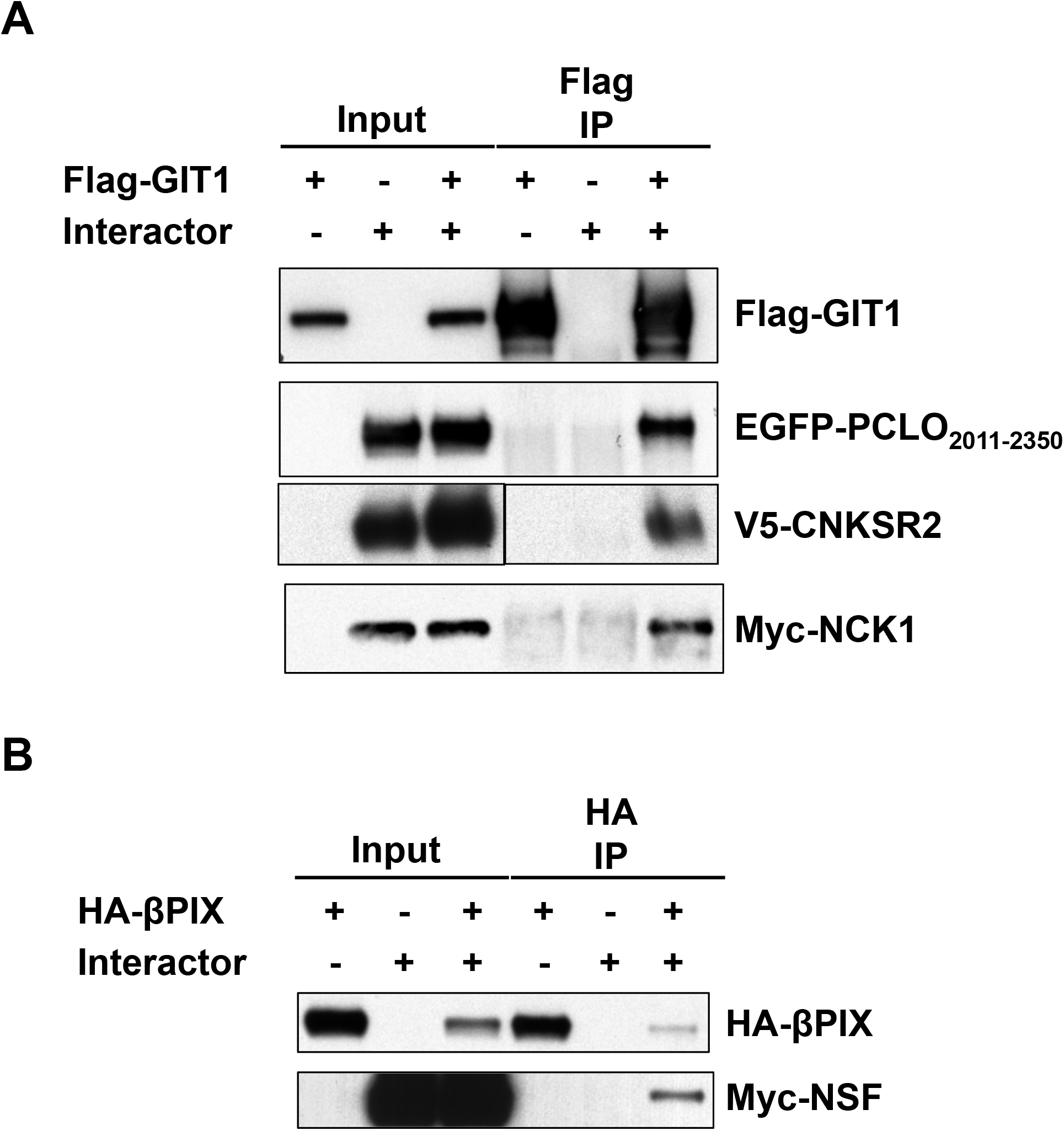
Confirmation of physical interactions between GIT1 and a set of GIT1-regulated phospho-proteins. Co-immunoprecipitation assays carried out in HEK293 cells grown in 6-well plates and transfected (using Lipofectamine 2000) with the indicated epitope-tagged versions of GIT1 and interacting proteins. Immunoprecipitations were performed using anti-Flag antibody (panel **A**) or anti-HA antibody (panel **B**) exactly as described in Kim et al., 2017. Detection of the co-immunoprecipitated interacting proteins was by western blotting with antibodies directed against the epitope tags of the interacting proteins.

**Supplemental Figure 3:**
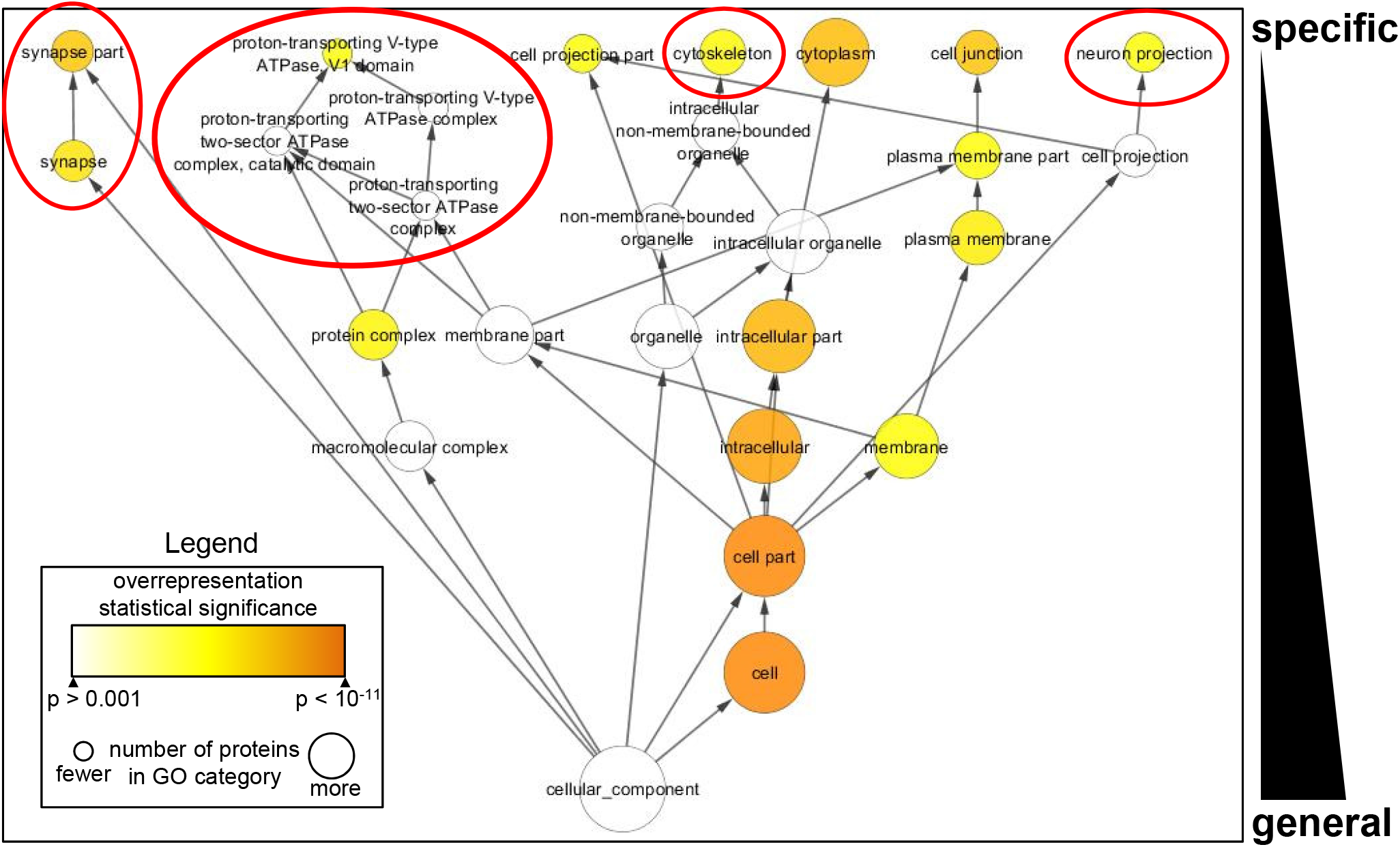
BiNGO visualization of GO cellular component category enrichment in robustly GIT1-regulated phospho-proteins. Global analysis of GO cellular component category enrichment (using a stringent criterion: p < 0.001) in these GIT1-regulated (≥ 2-fold) phospho-proteins identified a small network of components including several known to be associated with GIT1 (for a review see Zhou et al., 2016): cytoskeleton (proteins MACF1, ARHGAP21, CHRM4, STMN1, MAP1A, AKAP5, NEFH, SRCIN1, UBXN6, EIF3A, and SEPT3); neuron projection (proteins NSF, PAK1, SCN2A1, CHRM4, AKAP5, NEFH, and SRCIN1); and synapse (proteins RIMS1, CHRM4, GABRA4, AKAP5, SRCIN1, SYT7, SEPT3, and SHANK1); and one component not previously associated with GIT1 function: V-type ATPases (proteins ATP6V1H and ATP6V1C1). The enrichment of this last cellular component, V-type ATPases (which are critical components of synaptic vesicles; for a review see Morel & Poea-Guyon, 2015) is perhaps not surprising given GIT1’s known regulation of synaptic vesicles (for a review see Zhou et al., 2016).

## Supplemental Table Legends

**Supplemental Table 1: List of proteins detected in mouse hippocampus using TMT-based quantitative mass spectrometry.**

**Supplemental Table 2: List of GO categories enriched (PANTHER analysis) in proteins detected in mouse hippocampus using TMT-based quantitative mass spectrometry.**

**Supplemental Table 3: List of protein phosphorylation sites detected in mouse hippocampus using TMT-based quantitative mass spectrometry.**

**Supplemental Table 4: Previously known and novel phosphorylation sites detected on GIT1 and GRIN2A in mouse hippocampus using TMT-based quantitative mass spectrometry.**

**Supplemental Table 5: Statistically enriched GO cellular component categories (BINGO analysis) in the total set of phospho-proteins detected in mouse hippocampus using TMT-based quantitative mass spectrometry.**

**Supplemental Table 6: Proteins detected in mouse hippocampus using TMT-based quantitative mass spectrometry that are encoded within genomic loci with common variation associated with schizophrenia that contain only a single protein-coding gene.**

**Supplemental Table 7: List of normalized fold changes in protein phosphorylation levels induced by *GIT1* neural knockout in mouse hippocampus measured using TMT-based quantitative mass spectrometry.**

**Supplemental Table 8: Reactome database molecular pathways enriched among the set of GIT1-regulated phospho-proteins in mouse hippocampus measured using TMT-based quantitative mass spectrometry.**

**Supplemental Table 9: List of GO categories enriched (PANTHER analysis) in GIT1-regulated phospho-proteins detected in mouse hippocampus using TMT-based quantitative mass spectrometry.**

**Supplemental Table 10: List of GO categories enriched (PANTHER analysis) in GIT1-regulated phospho-proteins detected in mouse hippocampus using TMT-based quantitative mass spectrometry that describe a specific cellular compartment (shown in treemap in Figure 3B).**

**Supplemental Table 11: List of GIT1-regulated hippocampal phospho-proteins that have common genetic variants associated with schizophrenia.**

**Supplemental File 1: Cytoscape hierarchical network of statistically enriched GO cellular component categories found in the total set of all phospho-proteins detected in mouse hippocampus using TMT-based quantitative mass spectrometry.**

## References

Allen KM, Gleeson JG, Bagrodia S, Partington MW, MacMillan JC, Cerione RA, Mulley JC, Walsh CA. PAK3 mutation in nonsyndromic X-linked mental retardation. Nat Genet. 1998 Sep;20(1):25–30.

Aslan JE, Baker SM, Loren CP, Haley KM, Itakura A, Pang J, Greenberg DL, David LL, Manser E, Chernoff J, McCarty OJ. The PAK system links Rho GTPase signaling to thrombin-mediated platelet activation. Am J Physiol Cell Physiol. 2013 Sep;305(5):C519–28. doi: 10.1152/ajpcell.00418.2012. Epub 2013 Jun 19. PubMed PMID: 23784547; PubMed Central PMCID: PMC3761148.

Bagrodia S, Bailey D, Lenard Z, Hart M, Guan JL, Premont RT, Taylor SJ, Cerione RA. A tyrosine-phosphorylated protein that binds to an important regulatory region on the cool family of p21-activated kinase-binding proteins. J Biol Chem. 1999 Aug 6;274(32):22393–400.

Balu D, Larson JR, Schmidt JV, Wirtshafter D, Yevtodiyenko A, Leonard JP. Behavioral and physiological characterization of PKC-dependent phosphorylation in the Grin2aΔPKC mouse. Brain Res. 2016 Sep 1;1646:315–26.

Bates B, Rios M, Trumpp A, Chen C, Fan G, Bishop JM, Jaenisch R. Neurotrophin-3 is required for proper cerebellar development. Nat Neurosci. 1999 Feb;2(2):115–7. PubMed PMID: 10195193.

Boyken J, Grønborg M, Riedel D, Urlaub H, Jahn R, Chua JJ. Molecular profiling of synaptic vesicle docking sites reveals novel proteins but few differences between glutamatergic and GABAergic synapses. Neuron. 2013 Apr 24;78(2):285–97.

Braff D, Stone C, Callaway E, Geyer M, Glick I, Bali L. Prestimulus effects on human startle reflex in normals and schizophrenics. Psychophysiology. 1978Jul;15(4):339–43.

Broek JA, Lin Z, de Gruiter HM, van ’t Spijker H, Haasdijk ED, Cox D, Ozcan S, van Cappellen GW, Houtsmuller AB, Willemsen R, de Zeeuw CI, Bahn S. Synaptic vesicle dynamic changes in a model of fragile X. Mol Autism. 2016 Mar 1;7:17.

Collins MO, Husi H, Yu L, Brandon JM, Anderson CN, Blackstock WP, Choudhary JS, Grant SG. Molecular characterization and comparison of the components and multiprotein complexes in the postsynaptic proteome. J Neurochem. 2006 Apr;97 Suppl 1:16–23.

Cristino AS, Williams SM, Hawi Z, An JY, Bellgrove MA, Schwartz CE, Costa Lda F, Claudianos C. Neurodevelopmental and neuropsychiatric disorders represent an interconnected molecular system. Mol Psychiatry. 2014 Mar;19(3):294–301.

Datta D, Arion D, Corradi JP, Lewis DA. Altered expression of CDC42 signaling pathway components in cortical layer 3 pyramidal cells in schizophrenia. Biol Psychiatry. 2015 Dec 1;78(11):775–85.

Doherty JL, O’Donovan MC, Owen MJ. Recent genomic advances in schizophrenia. Clin Genet. 2012 Feb;81(2):103–9.

Dudchenko PA. An overview of the tasks used to test working memory in rodents. Neurosci Biobehav Rev. 2004 Nov;28(7):699–709.

Edwards RH. The neurotransmitter cycle and quantal size. Neuron. 2007 Sep 20;55(6):835–58. Review. PubMed PMID: 17880890.

Egbujo CN, Sinclair D, Hahn CG. Dysregulations of Synaptic Vesicle Trafficking in Schizophrenia. Curr Psychiatry Rep. 2016 Aug;18(8):77.

Elias JE, Gygi SP. 2010. Target-decoy search strategy for mass spectrometry-based proteomics. Methods in Molecular Biology 604: 55–71.

Endele S, Rosenberger G, Geider K, Popp B, Tamer C, Stefanova I, Milh M, Kortüm F, Fritsch A, Pientka FK, Hellenbroich Y, Kalscheuer VM, Kohlhase J, Moog U, Rappold G, Rauch A, Ropers HH, von Spiczak S, Tönnies H, Villeneuve N, Villard L, Zabel B, Zenker M, Laube B, Reis A, Wieczorek D, Van Maldergem L, Kutsche K. Mutations in GRIN2A and GRIN2B encoding regulatory subunits of NMDA receptors cause variable neurodevelopmental phenotypes. Nat Genet. 2010 Nov;42(11):1021–6.

Frese S, Schubert WD, Findeis AC, Marquardt T, Roske YS, Stradal TE, Heinz DW. The phosphotyrosine peptide binding specificity of Nck1 and Nck2 Src homology 2 domains. J Biol Chem. 2006 Jun 30;281(26):18236–45. Epub 2006 Apr 24. PubMed PMID: 16636066.

Fromer M, Pocklington AJ, Kavanagh DH, Williams HJ, Dwyer S, Gormley P, Georgieva L, Rees E, Palta P, Ruderfer DM, Carrera N, Humphreys I, Johnson JS, Roussos P, Barker DD, Banks E, Milanova V, Grant SG, Hannon E, Rose SA, Chambert K, Mahajan M, Scolnick EM, Moran JL, Kirov G, Palotie A, McCarroll SA, Holmans P, Sklar P, Owen MJ, Purcell SM, O’Donovan MC. De novo mutations in schizophrenia implicate synaptic networks. Nature. 2014 Feb 13;506(7487):179–84.

Fu CA, Shen M, Huang BC, Lasaga J, Payan DG, Luo Y. TNIK, a novel member of the germinal center kinase family that activates the c-Jun N-terminal kinase pathway and regulates the cytoskeleton. J Biol Chem. 1999 Oct22;274(43):30729–37. PubMed PMID: 10521462.

Genovese G, Fromer M, Stahl EA, Ruderfer DM, Chambert K, Landén M, Moran JL, Purcell SM, Sklar P, Sullivan PF, Hultman CM, McCarroll SA. Increased burden of ultra-rare protein-altering variants among 4,877 individuals with schizophrenia. Nat Neurosci. 2016 Oct 3.

González-Burgos I, Feria-Velasco A. Serotonin/dopamine interaction in memory formation. Prog Brain Res. 2008;172:603–23.

Hall J, Trent S, Thomas KL, O’Donovan MC, Owen MJ. Genetic risk for schizophrenia: convergence on synaptic pathways involved in plasticity. Biol Psychiatry. 2015 Jan 1;77(1):52–8.

Harrison PJ. The hippocampus in schizophrenia: a review of the neuropathological evidence and its pathophysiological implications. Psychopharmacology (Berl). 2004 Jun;174(1):151–62.

Hayashida M, Tanifuji S, Ma H, Murakami N, Mochida S. Neural activity selects myosin IIB and VI with a specific time window in distinct dynamin isoform-mediated synaptic vesicle reuse pathways. J Neurosci. 2015 Jun 10;35(23):8901–13.

Hormozdiari F, Penn O, Borenstein E, Eichler EE. The discovery of integrated gene networks for autism and related disorders. Genome Res. 2015 Jan;25(1):142–54.

Houge G, Rasmussen IH, Hovland R. Loss-of-Function CNKSR2 Mutation Is a Likely Cause of Non-Syndromic X-Linked Intellectual Disability. Mol Syndromol. 2012 Jan;2(2):60–63.

Hsu RM, Tsai MH, Hsieh YJ, Lyu PC, Yu JS. Identification of MYO18A as a novel interacting partner of the PAK2/betaPIX/GIT1 complex and its potential function in modulating epithelial cell migration. Mol Biol Cell. 2010 Jan 15;21(2):287–301. doi: 10.1091/mbc.E09-03-0232. Epub 2009 Nov 18. PubMed PMID:19923322; PubMed Central PMCID: PMC2808764.

Jaudon F, Raynaud F, Wehrlé R, Bellanger JM, Doulazmi M, Vodjdani G, Gasman S, Fagni L, Dusart I, Debant A, Schmidt S. The RhoGEF DOCK10 is essential for dendritic spine morphogenesis. Mol Biol Cell. 2015 Jun 1;26(11):2112–27.

Jordi E, Heiman M, Marion-Poll L, Guermonprez P, Cheng SK, Nairn AC, Greengard P, Girault JA. Differential effects of cocaine on histone posttranslational modifications in identified populations of striatal neurons. Proc Natl Acad Sci U S A. 2013 Jun 4;110(23):9511–6.

Ka M, Kim WY. Microtubule-Actin Crosslinking Factor 1 Is Required for Dendritic Arborization and Axon Outgrowth in the Developing Brain. Mol Neurobiol. 2016 Nov;53(9):6018–6032.

Kahn RS, Keefe RS. Schizophrenia is a cognitive illness: time for a change in focus. JAMA Psychiatry. 2013 Oct;70(10):1107–12.

Kim JJ, Fanselow MS. Modality-specific retrograde amnesia of fear. Science. 1992 May 1;256(5057):675–7.

Kim S, Ko J, Shin H, Lee JR, Lim C, Han JH, Altrock WD, Garner CC, Gundelfinger ED, Premont RT, Kaang BK, Kim E. The GIT family of proteins forms multimers and associates with the presynaptic cytomatrix protein Piccolo. J Biol Chem. 2003 Feb 21;278(8):6291–300.

Kim K, Lakhanpal G, Lu HE, Khan M, Suzuki A, Hayashi MK, Narayanan R, Luyben TT, Matsuda T, Nagai T, Blanpied TA, Hayashi Y, Okamoto K. A Temporary Gating of Actin Remodeling during Synaptic Plasticity Consists of the Interplay between the Kinase and Structural Functions of CaMKII. Neuron. 2015 Aug 19;87(4):813–26. doi:10.1016/j.neuron.2015.07.023. Erratum in: Neuron. 2015 Oct 21;88(2):433. PubMed PMID: 26291163; PubMed Central PMCID: PMC4548268.

Kim MJ, Biag J, Fass DM, Lewis MC, Zhang Q, Fleishman M, Gangwar SP, Machius M, Fromer M, Purcell SM, McCarroll SA, Rudenko G, Premont RT, Scolnick EM, Haggarty SJ. Functional analysis of rare variants found in schizophrenia implicates a critical role for GIT1-PAK3 signaling in neuroplasticity. Mol Psychiatry. 2017 Mar;22(3):417–429. doi: 10.1038/mp.2016.98. Epub 2016 Jul 26. PubMed PMID: 27457813.

Klemmer P, Smit AB, Li KW. Proteomics analysis of immuno-precipitated synaptic protein complexes. J Proteomics. 2009 Feb 15;72(1):82–90.

Ko J, Kim S, Valtschanoff JG, Shin H, Lee JR, Sheng M, Premont RT, Weinberg RJ, Kim E. Interaction between liprin-alpha and GIT1 is required for AMPA receptor targeting. J Neurosci. 2003 Mar 1;23(5):1667–77.

Ko J, Na M, Kim S, Lee JR, Kim E. Interaction of the ERC family of RIM-binding proteins with the liprin-alpha family of multidomain proteins. J Biol Chem. 2003b Oct 24;278(43):42377–85. Epub 2003 Aug 15. PubMed PMID: 12923177.

Kohansal-Nodehi M, Chua JJ, Urlaub H, Jahn R, Czernik D. Analysis of protein phosphorylation in nerve terminal reveals extensive changes in active zone proteins upon exocytosis. Elife. 2016 Apr 26;5. pii: e14530. doi:10.7554/eLife.14530. PubMed PMID: 27115346; PubMed Central PMCID: PMC4894758.

Kutsche K, Yntema H, Brandt A, Jantke I, Nothwang HG, Orth U, Boavida MG, David D, Chelly J, Fryns JP, Moraine C, Ropers HH, Hamel BC, van Bokhoven H, Gal A. Mutations in ARHGEF6, encoding a guanine nucleotide exchange factor for Rho GTPases, in patients with X-linked mental retardation. Nat Genet. 2000 Oct;26(2):247–50.

Lalonde R. The neurobiological basis of spontaneous alternation. Neurosci Biobehav Rev. 2002 Jan;26(1):91–104.

Lek M, Karczewski KJ, Minikel EV, Samocha KE, Banks E, Fennell T, O’Donnell-Luria AH, Ware JS, Hill AJ, Cummings BB, Tukiainen T, Birnbaum DP, Kosmicki JA, Duncan LE, Estrada K, Zhao F, Zou J, Pierce-Hoffman E, Berghout J, Cooper DN, Deflaux N, DePristo M, Do R, Flannick J, Fromer M, Gauthier L, Goldstein J, Gupta N, Howrigan D, Kiezun A, Kurki MI, Moonshine AL, Natarajan P, Orozco L, Peloso GM, Poplin R, Rivas MA, Ruano-Rubio V, Rose SA, Ruderfer DM, Shakir K, Stenson PD, Stevens C, Thomas BP, Tiao G, Tusie-Luna MT, Weisburd B, Won HH, Yu D, Altshuler DM, Ardissino D, Boehnke M, Danesh J, Donnelly S, Elosua R, Florez JC, Gabriel SB, Getz G, Glatt SJ, Hultman CM, Kathiresan S, Laakso M, McCarroll S, McCarthy MI, McGovern D, McPherson R, Neale BM, Palotie A, Purcell SM, Saleheen D, Scharf JM, Sklar P, Sullivan PF, Tuomilehto J, Tsuang MT, Watkins HC, Wilson JG, Daly MJ, MacArthur DG; Exome Aggregation Consortium. Analysis of protein-coding genetic variation in 60,706 humans. Nature. 2016 Aug 17;536(7616):285–91.

Lim J, Ritt DA, Zhou M, Morrison DK. The CNK2 scaffold interacts with vilse and modulates Rac cycling during spine morphogenesis in hippocampal neurons. Curr Biol. 2014 Mar 31;24(7):786–92.

Maere S, Heymans K, Kuiper M. BiNGO: a Cytoscape plugin to assess overrepresentation of gene ontology categories in biological networks. Bioinformatics. 2005 Aug 15;21(16):3448–9. Epub 2005 Jun 21. PubMed PMID:15972284.

Martin HG, Henley JM, Meyer G. Novel putative targets of N-ethylmaleimide sensitive fusion protein (NSF) and alpha/beta soluble NSF attachment proteins (SNAPs) include the Pak-binding nucleotide exchange factor betaPIX. J Cell Biochem. 2006 Nov 1;99(4):1203–15. PubMed PMID: 16795052; PubMed Central PMCID:PMC3308139.

Martyn AC, Toth K, Schmalzigaug R, Hedrick NG, Rodriguiz RM, Yasuda R, Wetsel, WC, & Premont RT. GIT1 regulates synaptic structural plasticity underlying learning. PLOS ONE. 2018 epub March 19, 2018 https://doi.org/10.1371/journal.pone.0194350

McGrath J, Saha S, Chant D, Welham J. Schizophrenia: a concise overview of incidence, prevalence, and mortality. Epidemiol Rev. 2008;30:67–76.

McRae et al., Deciphering Developmental Disorders Study. Prevalence and architecture of de novo mutations in developmental disorders. Nature. 2017 Feb 23;542(7642):433–438. doi: 10.1038/nature21062. Epub 2017 Jan 25. PubMed PMID: 28135719.

Menon P, Deane R, Sagare A, Lane SM, Zarcone TJ, O’Dell MR, Yan C, Zlokovic BV, Berk BC. Impaired spine formation and learning in GPCR kinase 2 interacting protein-1 (GIT1) knockout mice. Brain Res. 2010 Mar 4;1317:218–26.

Mertins P, Mani DR, Ruggles KV, Gillette MA, Clauser KR, Wang P, Wang X, Qiao JW, Cao S, Petralia F, Kawaler E, Mundt F, Krug K, Tu Z, Lei JT, Gatza ML, Wilkerson M, Perou CM, Yellapantula V, Huang KL, Lin C, McLellan MD, Yan P, Davies SR, Townsend RR, Skates SJ, Wang J, Zhang B, Kinsinger CR, Mesri M, Rodriguez H, Ding L, Paulovich AG, Fenyö D, Ellis MJ, Carr SA; NCI CPTAC. Proteogenomics connects somatic mutations to signalling in breast cancer. Nature. 2016 Jun 2;534(7605):55–62. doi: 10.1038/nature18003. Epub 2016 May 25. PubMed PMID: 27251275; PubMed Central PMCID: PMC5102256.

Montesinos MS, Dong W, Goff K, Das B, Guerrero-Given D, Schmalzigaug R, Premont RT, Satterfield R, Kamasawa N, Young SM Jr. Presynaptic Deletion of GIT Proteins Results in Increased Synaptic Strength at a Mammalian Central Synapse. Neuron. 2015 Dec 2;88(5):918–25.

Morel N, Poëa-Guyon S. The membrane domain of vacuolar H(+)ATPase: a crucial player in neurotransmitter exocytotic release. Cell Mol Life Sci. 2015 Jul;72(13):2561–73.

Muller J, Corodimas KP, Fridel Z, LeDoux JE. Functional inactivation of the lateral and basal nuclei of the amygdala by muscimol infusion prevents fear conditioning to an explicit conditioned stimulus and to contextual stimuli. Behav Neurosci. 1997 Aug;111(4):683–91.

Mundt F, Rajput S, Li S, Ruggles KV, Mooradian AD, Mertins P, Gillette MA, Krug K, Guo Z, Hoog J, Erdmann-Gilmore P, Primeau T, Huang S, Edwards DP, Wang X, Wang X, Kawaler E, Mani DR, Clauser KR, Gao F, Luo J, Davies SR, Johnson GL, Huang KL, Yoon CJ, Ding L, Fenyö D, Ellis MJ, Townsend RR, Held JM, Carr SA, Ma CX. Mass spectrometry-based proteomics reveals potential roles of NEK9 and MAP2K4 in resistance to PI3K inhibitors in triple negative breast cancers. Cancer Res. 2018 Feb 22. pii: canres.1990.2017. doi: 10.1158/0008-5472.CAN-17-1990. [Epub ahead of print] PubMed PMID: 29472518.

Nesvizhskii AI, Aebersold R. 2005. Interpretation of shotgun proteomic data - The protein inference problem. Molecular & Cellular Proteomics 4(10): 1419–1440.

Pang J, Yan C, Natarajan K, Cavet ME, Massett MP, Yin G, Berk BC. GIT1 mediates HDAC5 activation by angiotensin II in vascular smooth muscle cells. Arterioscler Thromb Vasc Biol. 2008 May;28(5):892–8. doi:10.1161/ATVBAHA.107.161349. Epub 2008 Feb 21. PubMed PMID: 18292392; PubMedCentral PMCID: PMC2735338.

Pang J, Hoefen R, Pryhuber GS, Wang J, Yin G, White RJ, Xu X, O’Dell MR, Mohan A, Michaloski H, Massett MP, Yan C, Berk BC. G-protein-coupled receptor kinase interacting protein-1 is required for pulmonary vascular development. Circulation. 2009 Mar 24;119(11):1524–32.

Pardiñas AF, Holmans P, Pocklington AJ, Escott-Price V, Ripke S, Carrera N, Legge SE, Bishop S, Cameron D, Hamshere ML, Han J, Hubbard L, Lynham A, Mantripragada K, Rees E, MacCabe JH, McCarroll SA, Baune BT, Breen G, Byrne EM, Dannlowski U, Eley TC, Hayward C, Martin NG, McIntosh AM, Plomin R, Porteous DJ, Wray NR, Caballero A, Geschwind DH, Huckins LM, Ruderfer DM, Santiago E, Sklar P, Stahl EA, Won H, Agerbo E, Als TD, Andreassen OA, Bækvad-Hansen M, Mortensen PB, Pedersen CB, Børglum AD, Bybjerg-Grauholm J, Djurovic S, Durmishi N, Pedersen MG, Golimbet V, Grove J, Hougaard DM, Mattheisen M, Molden E, Mors O, Nordentoft M, Pejovic-Milovancevic M, Sigurdsson E, Silagadze T, Hansen CS, Stefansson K, Stefansson H, Steinberg S, Tosato S, Werge T; GERAD1 Consortium:; CRESTAR Consortium:, Collier DA, Rujescu D, Kirov G, Owen MJ, O’Donovan MC, Walters JTR; GERAD1 Consortium; CRESTAR Consortium; GERAD1 Consortium; CRESTAR Consortium. Common schizophrenia alleles are enriched in mutation-intolerant genes and in regions under strong background selection. Nat Genet. 2018 Feb 26. doi:10.1038/s41588-018-0059-2. [Epub ahead of print] PubMed PMID: 29483656.

Park E, Na M, Choi J, Kim S, Lee JR, Yoon J, Park D, Sheng M, Kim E. The Shank family of postsynaptic density proteins interacts with and promotes synaptic accumulation of the beta PIX guanine nucleotide exchange factor for Rac1 and Cdc42. J Biol Chem. 2003 May 23;278(21):19220–9. Epub 2003 Mar 7. PubMed PMID:12626503.

Pechstein A, Shupliakov O, Haucke V. Intersectin 1: a versatile actor in the synaptic vesicle cycle. Biochem Soc Trans. 2010 Feb;38(Pt 1):181–6.

Purcell SM, Moran JL, Fromer M, Ruderfer D, Solovieff N, Roussos P, O’Dushlaine C, Chambert K, Bergen SE, Kähler A, Duncan L, Stahl E, Genovese G, Fernández E, Collins MO, Komiyama NH, Choudhary JS, Magnusson PK, Banks E, Shakir K, Garimella K, Fennell T, DePristo M, Grant SG, Haggarty SJ, Gabriel S, Scolnick EM, Lander ES, Hultman CM, Sullivan PF, McCarroll SA, Sklar P. A polygenic burden of rare disruptive mutations in schizophrenia. Nature. 2014 Feb13;506(7487):185–90.

Rennefahrt UE, Deacon SW, Parker SA, Devarajan K, Beeser A, Chernoff J, Knapp S, Turk BE, Peterson JR. Specificity profiling of Pak kinases allows identification of novel phosphorylation sites. J Biol Chem. 2007 May 25;282(21):15667–78. Epub 2007 Mar 28. PubMed PMID: 17392278.

Sanders SJ, He X, Willsey AJ, Ercan-Sencicek AG, Samocha KE, Cicek AE, Murtha MT, Bal VH, Bishop SL, Dong S, Goldberg AP, Jinlu C, Keaney JF 3rd, Klei L, Mandell JD, Moreno-De- Luca D, Poultney CS, Robinson EB, Smith L, Solli-Nowlan T, Su MY, Teran NA, Walker MF, Werling DM, Beaudet AL, Cantor RM, Fombonne E, Geschwind DH, Grice DE, Lord C, Lowe JK, Mane SM, Martin DM, Morrow EM, Talkowski ME, Sutcliffe JS, Walsh CA, Yu TW; Autism Sequencing Consortium, Ledbetter DH, Martin CL, Cook EH, Buxbaum JD, Daly MJ, Devlin B, Roeder K, State MW. Insights into Autism Spectrum Disorder Genomic Architecture and Biology from 71 Risk Loci. Neuron. 2015 Sep 23;87(6):1215–1233. doi: 10.1016/j.neuron.2015.09.016. PubMed PMID: 26402605; PubMed Central PMCID: PMC4624267.

Schizophrenia Working Group of the Psychiatric Genomics Consortium. Biological insights from 108 schizophrenia-associated genetic loci. Nature. 2014 Jul 24;511(7510):421–7.

Schmalzigaug R, Rodriguiz RM, Bonner PE, Davidson CE, Wetsel WC, Premont RT. Impaired fear response in mice lacking GIT1. Neurosci Lett. 2009 Jul 17;458(2):79–83.

Shin H, Wyszynski M, Huh KH, Valtschanoff JG, Lee JR, Ko J, Streuli M, Weinberg RJ, Sheng M, Kim E. Association of the kinesin motor KIF1A with the multimodular protein liprin-alpha. J Biol Chem. 2003 Mar 28;278(13):11393–401. Epub 2003 Jan 8. PubMed PMID: 12522103.

Shin EY, Shim ES, Lee CS, Kim HK, Kim EG. Phosphorylation of RhoGDI1 by p21-activated kinase 2 mediates basic fibroblast growth factor-stimulated neurite outgrowth in PC12 cells. Biochem Biophys Res Commun. 2009 Feb 6;379(2):384–9. doi: 10.1016/j.bbrc.2008.12.066. Epub 2008 Dec 25. PubMed PMID: 19103160.

Silverman JL, Yang M, Lord C, Crawley JN. Behavioural phenotyping assays for mouse models of autism. Nat Rev Neurosci. 2010 Jul;11(7):490–502. doi:10.1038/nrn2851. Review. PubMed PMID: 20559336; PubMed Central PMCID: PMC3087436.

Südhof TC. The presynaptic active zone. Neuron. 2012 Jul 12;75(1):11–25.

Tandon R, Gaebel W, Barch DM, Bustillo J, Gur RE, Heckers S, Malaspina D, Owen MJ, Schultz S, Tsuang M, Van Os J, Carpenter W. Definition and description of schizophrenia in the DSM-5. Schizophr Res. 2013 Oct;150(1):3–10.

Uddin M, Pellecchia G, Thiruvahindrapuram B, D’Abate L, Merico D, Chan A, Zarrei M, Tammimies K, Walker S, Gazzellone MJ, Nalpathamkalam T, Yuen RK, Devriendt K, Mathonnet G, Lemyre E, Nizard S, Shago M, Joseph-George AM, Noor A, Carter MT, Yoon G, Kannu P, Tihy F, Thorland EC, Marshall CR, Buchanan JA, Speevak M, Stavropoulos DJ, Scherer SW. Indexing Effects of Copy Number Variation on Genes Involved in Developmental Delay. Sci Rep. 2016 Jul 1;6:28663.

Vadlamudi RK, Manavathi B, Singh RR, Nguyen D, Li F, Kumar R. An essential role of Pak1 phosphorylation of SHARP in Notch signaling. Oncogene. 2005 Jun 30;24(28):4591–6. PubMed PMID: 15824732.

Veeranna, Amin ND, Ahn NG, Jaffe H, Winters CA, Grant P, Pant HC. Mitogen-activated protein kinases (Erk1,2) phosphorylate Lys-Ser-Pro (KSP) repeats in neurofilament proteins NF-H and NF-M. J Neurosci. 1998 Jun 1;18(11):4008–21. PubMed PMID: 9592082.

Versele M, Thorner J. Septin collar formation in budding yeast requires GTP binding and direct phosphorylation by the PAK, Cla4. J Cell Biol. 2004 Mar1;164(5):701–15. PubMed PMID: 14993234; PubMed Central PMCID: PMC2172161.

Vizi ES, Kiss JP. Neurochemistry and pharmacology of the major hippocampal transmitter systems: synaptic and nonsynaptic interactions. Hippocampus.1998;8(6):566–607.

Vissers LE, Gilissen C, Veltman JA. Genetic studies in intellectual disability and related disorders. Nat Rev Genet. 2016 Jan;17(1):9–18.

Volk L, Chiu SL, Sharma K, Huganir RL. Glutamate synapses in human cognitive disorders. Annu Rev Neurosci. 2015 Jul 8;38:127–49.

Waites CL, Leal-Ortiz SA, Andlauer TF, Sigrist SJ, Garner CC. Piccolo regulates the dynamic assembly of presynaptic F-actin. J Neurosci. 2011 Oct 5;31(40):14250–63.

Wang Q, Amato SP, Rubitski DM, Hayward MM, Kormos BL, Verhoest PR, Xu L, Brandon NJ, Ehlers MD. Identification of Phosphorylation Consensus Sequences and Endogenous Neuronal Substrates of the Psychiatric Risk Kinase TNIK. J Pharmacol Exp Ther. 2016 Feb;356(2):410–23. doi: 10.1124/jpet.115.229880. Epub 2015 Dec 8. PubMed PMID: 26645429.

Watanabe M, Nomura K, Ohyama A, Ishikawa R, Komiya Y, Hosaka K, Yamauchi E, Taniguchi H, Sasakawa N, Kumakura K, Ushiki T, Sato O, Ikebe M, Igarashi M. Myosin-Va regulates exocytosis through the submicromolar Ca2+-dependent binding of syntaxin-1A. Mol Biol Cell. 2005 Oct;16(10):4519–30.

Wittmann T, Bokoch GM, Waterman-Storer CM. Regulation of microtubule destabilizing activity of Op18/stathmin downstream of Rac1. J Biol Chem. 2004 Feb 13;279(7):6196–203. Epub 2003 Nov 26. PubMed PMID: 14645234.

Won H, Mah W, Kim E, Kim JW, Hahm EK, Kim MH, Cho S, Kim J, Jang H, Cho SC, Kim BN, Shin MS, Seo J, Jeong J, Choi SY, Kim D, Kang C, Kim E. GIT1 is associated with ADHD in humans and ADHD-like behaviors in mice. Nat Med. 2011 May;17(5):566–72.

Woolfrey KM, Dell’Acqua ML. Coordination of Protein Phosphorylation and Dephosphorylation in Synaptic Plasticity. J Biol Chem. 2015 Nov 27;290(48):28604–12. doi:10.1074/jbc.R115.657262.

Xue J, Tsang CW, Gai WP, Malladi CS, Trimble WS, Rostas JA, Robinson PJ. Septin 3 (G-septin) is a developmentally regulated phosphoprotein enriched in presynaptic nerve terminals. J Neurochem. 2004 Nov;91(3):579–90.

Yao I, Hata Y, Ide N, Hirao K, Deguchi M, Nishioka H, Mizoguchi A, Takai Y. MAGUIN, a novel neuronal membrane-associated guanylate kinase-interacting protein. J Biol Chem. 1999 Apr 23;274(17):11889–96.

Yin G, Haendeler J, Yan C, Berk BC. GIT1 functions as a scaffold for MEK1-extracellular signal-regulated kinase 1 and 2 activation by angiotensin II and epidermal growth factor. Mol Cell Biol. 2004 Jan;24(2):875–85. PubMed PMID:14701758; PubMed Central PMCID: PMC343801.

Yin G, Zheng Q, Yan C, Berk BC. GIT1 is a scaffold for ERK1/2 activation in focal adhesions. J Biol Chem. 2005 Jul 29;280(30):27705–12. Epub 2005 May 27. PubMed PMID: 15923189.

Zhang H, Webb DJ, Asmussen H, Horwitz AF. Synapse formation is regulated by the signaling adaptor GIT1. J Cell Biol. 2003 Apr 14;161(1):131–42.

Zhang H, Webb DJ, Asmussen H, Niu S, Horwitz AF. A GIT1/PIX/Rac/PAK signaling module regulates spine morphogenesis and synapse formation through MLC. J Neurosci. 2005 Mar 30;25(13):3379–88.

Zhang N, Cai W, Yin G, Nagel DJ, Berk BC. GIT1 is a novel MEK1-ERK1/2 scaffold that localizes to focal adhesions. Cell Biol Int. 2009 Dec 16;34(1):41–7. doi:10.1042/CBI20090016. PubMed PMID: 19947948; PubMed Central PMCID: PMC3125965.

Zhao ZS, Manser E, Loo TH, Lim L. Coupling of PAK-interacting exchange factor PIX to GIT1 promotes focal complex disassembly. Mol Cell Biol. 2000 Sep;20(17):6354–63. PubMed PMID: 10938112; PubMed Central PMCID: PMC86110.

Zhou W, Li X, Premont RT. Expanding functions of GIT Arf GTPase-activating proteins, PIX Rho guanine nucleotide exchange factors and GIT-PIX complexes. J Cell Sci. 2016 May 15;129(10):1963–74.

