## Supplementary Materials for "Brain-Specific Deletion of GIT1 Impairs Cognition and Alters Phosphorylation of Synaptic Protein Networks Implicated in Schizophrenia Susceptibility"

### Slide 1
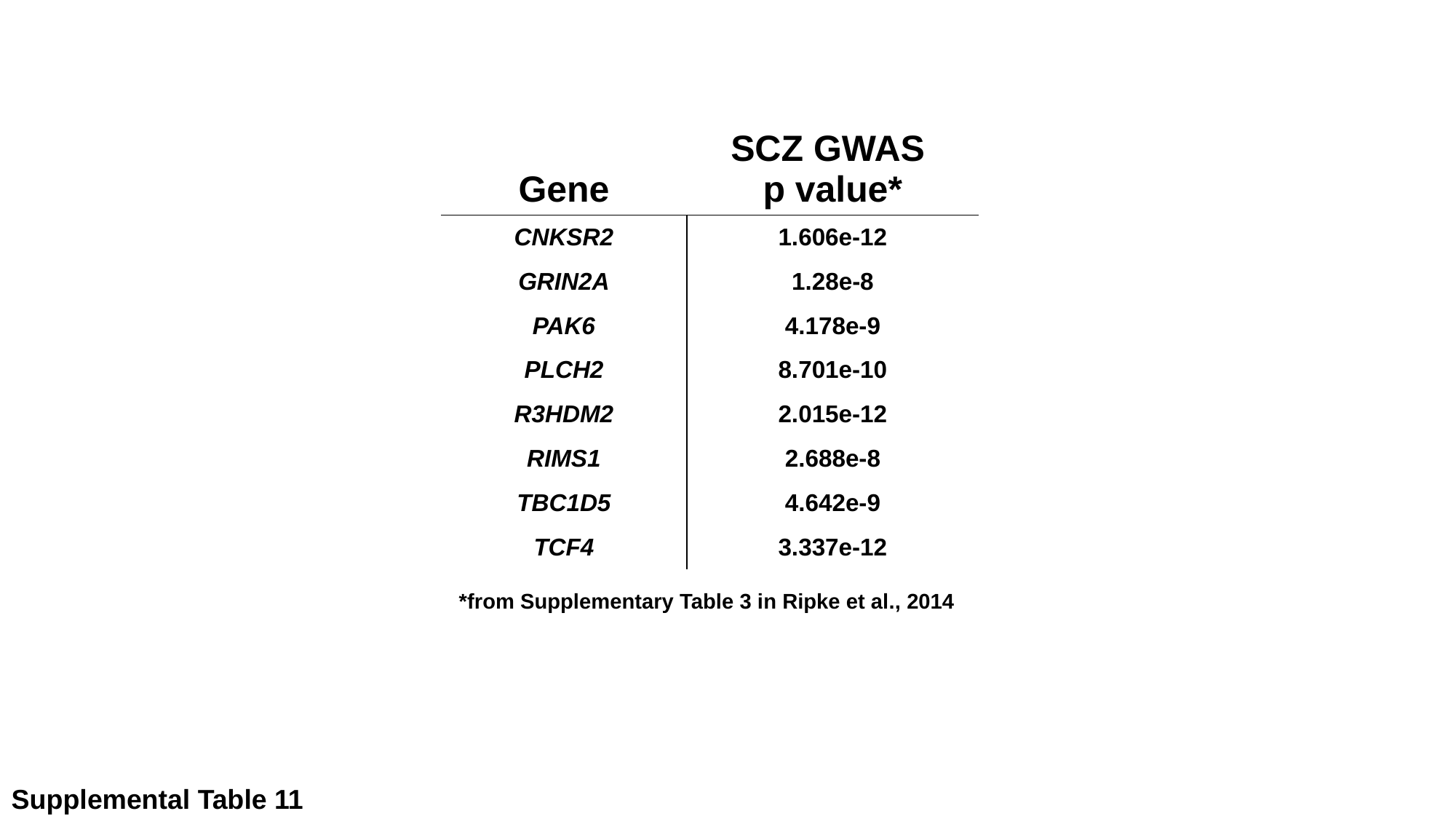

| Gene | SCZ GWAS p value\* |
| --- | --- |
| CNKSR2 | 1.606e-12 |
| Grin2a | 1.28e-8 |
| Pak6 | 4.178e-9 |
| Plch2 | 8.701e-10 |
| R3hdm2 | 2.015e-12 |
| Rims1 | 2.688e-8 |
| Tbc1d5 | 4.642e-9 |
| TCF4 | 3.337e-12 |
*from Supplementary Table 3 in Ripke et al., 2014
Supplemental Table 11
